# Characterising semantic prioritisation in visual working memory

**DOI:** 10.64898/2026.02.20.706943

**Authors:** Casper Kerrén, Carlos González-García, Juan Linde-Domingo

## Abstract

Visual working memory preserves multiple features of recently encountered information, but these features are not necessarily equally available to guide upcoming behaviour. In particular, categorical semantic content can be prioritised over perceptual content when objects are retrieved from memory. Here, we asked what drives this semantic prioritisation in visual working memory, and whether it depends on the attentional state of the remembered information. Across a reanalysis of a previous dataset and two preregistered experiments, participants showed a prioritisation of semantic-category information over perceptual one. Drift-diffusion modelling suggested that such an advantage arose from multiple components of decision formation, including processes beyond evidence accumulation. Critically, when a retro-cue allowed the relevant memory item to be prioritised before the probe, the semantic advantage in processes outside evidence accumulation almost disappeared, whereas its advantage during evidence accumulation remained largely unchanged. In a separate experiment, extending the retention interval alone did not reliably increase semantic prioritisation, while an intervening task produced a larger absolute advantage in pre- accumulation processing. These findings show that semantic prioritisation is not simply a fixed advantage of semantic judgements and that the relative availability of meaning and perceptual format in working memory is dynamically shaped by attentional prioritisation.

## Introduction

Adaptive behaviour depends on the ability to use information that is no longer available by current sensory input. To achieve this, cognitive systems must sustain internal representations over short intervals and make them available for ongoing computations. This capacity, known as working memory (WM), enables the use of task-relevant information beyond the moment of perception (Baddeley, 1992, 2012; D’Esposito & Postle, 2015). Despite its centrality, it remains unclear what format information takes in WM, how it is accessed, and to what extent WM representations resemble those instantiated during perception or those reconstructed from long-term memory (LTM).

A large body of research has investigated WM using delay-based paradigms, often involving the maintenance of a single stimulus or feature (D’Esposito & Postle, 2015; Pasternak & Greenlee, 2005). These studies have demonstrated that information held in WM can be decoded from brain sensory cortical regions typically associated with perceptual processing (Harrison & Tong, 2009; Serences et al., 2009), motivating influential sensory recruitment accounts. From this perspective, WM maintenance involves the continued engagement of perceptual representational formats. At the same time, converging behavioural and neural evidence indicates that WM representations are frequently transformed relative to their perceptual origin (Albers et al., 2013; Rademaker et al., 2019; Spaak et al., 2017). Rather than preserving a veridical sensory trace, WM may prioritise task-relevant dimensions, abstracted and semantic stimulus features (Kwak & Curtis, 2022; Pettini et al., 2025), or anticipated actions, depending on behavioural and attentional demands (Cipriani et al., 2024; Linde-Domingo & Spitzer, 2024; Myers et al., 2017).

Growing evidence indicates that taxonomic semantic knowledge can shape visual WM at multiple stages of processing. Meaningful real-world objects can be maintained more effectively than simple or unfamiliar stimuli (Brady et al., 2016), while semantic category coherence and functional relationships among objects can facilitate memory for their identities and locations (Hu & Jacobs, 2021; O’Donnell et al., 2018). More recent work has shown that graded semantic associations between objects predict visual working-memory performance independently of visual similarity, particularly when encoding time is limited (Yu et al., 2025). These findings suggest that more abstract knowledge can contribute to the organisation and accessibility of working-memory contents, although the processes underlying these benefits may differ across tasks and stages of processing.

Beyond the maintenance of a single item, WM is generally assumed to support the concurrent retention of multiple representational layers (Cowan, 2001; Jensen & Lisman, 1996; Miller, 1956). However, only some of these can occupy the focus of attention at any given moment (Cowan, 2001; Oberauer, 2002; Olivers et al., 2011). How information outside this focus is retained and accessed remains a matter of debate. Contemporary proposals range from transient synaptic traces that persist without sustained neural firing (Mongillo et al., 2008; Stokes, 2015), to distributed storage across cortical hierarchies at varying levels of abstraction. Others hold that unattended WM contents may rely on mechanisms closely related to those supporting long-term memory, particularly when they must be reactivated after a period of reduced attentional priority (Beukers et al., 2021; Jeneson & Squire, 2012). Recent theoretical and empirical work has extended this idea, suggesting that activity-silent WM may share computational properties with episodic memory retrieval (Beukers et al., 2021; Rose et al., 2016). From this perspective, whether information remains within the focus of attention, rather than how long it has been stored, may be a critical determinant of how it is accessed and used.

Insights from the episodic memory literature highlight a critical distinction between how information is accessed from memory and how it is processed during perception. During visual perception, neural processing typically unfolds along a feedforward trajectory in which low- level sensory attributes are generally resolved before higher-level, conceptual properties (Carlson et al., 2013; Cichy et al., 2014; Clarke & Tyler, 2015; DiCarlo et al., 2012; Serre et al., 2007). In contrast, during episodic retrieval, categorical conceptual or semantic information often becomes available more rapidly than categorical perceptual information (Lifanov et al., 2021; Linde-Domingo et al., 2019). This inversion of temporal dynamics has been interpreted as evidence that memory retrieval is reconstructive, potentially initiated by pattern completion mechanisms that reinstate cortical representations in a top-down manner (Kerrén et al., 2024, 2025, 2026; Norman & O’Reilly, 2003). Comparable temporal reversal patterns have also been observed in visual imagery (Dijkstra et al., 2018, 2020), reinforcing the idea that internally generated representations differ fundamentally from perceptually driven ones.

We previously asked whether similar access dynamics characterise WM. In Kerrén et al., (2022), we reported a robust prioritisation of categorical semantic over perceptual information when multiple items were maintained in working memory. Participants were faster, and often more accurate, in reporting semantic category than visuoperceptual format. However, in that study temporal distance was necessarily confounded with attentional allocation, leaving open whether lag itself, or the likelihood of attentional disengagement, drove the effect. These response dynamics resembled those observed during episodic memory retrieval (Lifanov et al., 2021; Linde-Domingo et al., 2019) and contrasted with those typically found in visual perception, where lower-level visual features are accessed earlier than higher-level conceptual attributes.

In these experiments, semantic information was operationalised as a binary category distinction (animal versus object), whereas perceptual information was operationalised as stimulus format (photograph versus line drawing). This approach allowed us to compare access to two predefined object dimensions using the same stimulus material, but did not assess richer semantic relationships, associative structure, or graded conceptual similarity. The observed prioritisation therefore concerns categorical semantic information rather than semantic knowledge in its broader sense.

Despite our previous findings, the processes underlying the prioritisation of categorical semantic over perceptual-format information in visual working memory remain unclear. Previous behavioural studies have demonstrated that semantic knowledge and associative relationships can facilitate working-memory performance (Hu & Jacobs, 2021; O’Donnell et al., 2018; Yu et al., 2025), while neural decoding studies have identified both perceptual and more abstract representational formats during maintenance (Liu et al., 2020). Although these approaches provide important insights into the content and fidelity of working-memory representations, they do not directly establish which stages of subsequent decision formation account for differences in the relative accessibility of semantic and perceptual information (van Ede & Nobre, 2023). One possibility is that the observed semantic advantage reflects processes preceding evidence accumulation, including the selection or reactivation of unattended representations. Alternatively, it may arise from differences in how semantic and perceptual information contributes to evidence accumulation or from decision-related biases, such as different response strategies (Griffin & Nobre, 2003). More generally, it remains unclear whether semantic prioritisation is driven primarily by attentional disruption during maintenance, the need to reinstate information from an unattended state, or the passage of time itself. Recent work further demonstrates that retrospective attention can preferentially enhance perceptual over semantic information (Cipriani et al., 2024), suggesting that attentional selection may influence different stages of information processing and decision formation.

In the present study, we build directly on these earlier findings to examine the conditions under which semantic prioritisation emerges in working memory. In particular, here we tested whether such bias in working memory arises prior to evidence accumulation or during the decision process itself. To adjudicate between access and accumulation-based accounts of semantic prioritisation, we applied drift diffusion modelling (DDM) to dissociate processes outside evidence accumulation (Ratcliff, 1978; Ratcliff & McKoon, 2008; Wiecki et al., 2013). First, we reanalysed our previous results (Kerrén et al., 2022) using DDM. Building on these results, in Experiment 1, we combined DDM with a retro-cue manipulation to assess whether this semantic preference is amplified when attentional selection is prevented and multiple items must remain concurrently maintained (Griffin & Nobre, 2003). In Experiment 2, we independently manipulated interference and delay during the maintenance interval to dissociate the effects of attentional disruption and time on the accessibility of semantic and perceptual information (Portrat et al., 2008; Ricker et al., 2010). Together, these experiments allow us to examine how semantic prioritisation is expressed through distinct components of decision formation and whether their relative contributions depend on attentional prioritisation, interference and retention time. By combining experimental manipulations with hierarchical drift-diffusion modelling, we aim to constrain the processes through which categorical semantic and perceptual-format information guide working-memory decisions, while recognising that these decision components do not uniquely identify the underlying retrieval or representational mechanisms.

## Results

### Drift-diffusion modelling of Kerrén et al (2022)

In Kerrén et al. (2022), we asked whether the semantic prioritisation observed during episodic memory retrieval (Linde-Domingo et al. 2019) also characterises visual working memory (VWM). Across two experiments, we observed a robust semantic advantage when participants were required to maintain more than one item simultaneously. This semantic prioritisation was systematically modulated by both memory load (the number of items studied, ranging from one to four) and the temporal interval between study and test (hereafter referred to as lag).

Importantly, this prioritisation of semantic over perceptual information is not a trivial consequence of stimulus properties (Fig. 1A). In a previous publication, using the same material and feature space, perceptual information (line-drawing vs. photograph) was accessed earlier than semantic features (animacy) when judgements were made during visual processing, without a memory demand (Linde-Domingo et al., 2019). This reversal suggests that the relative accessibility of perceptual and semantic features depends on whether processing is perceptually driven or internally generated.

**Figure 1.**
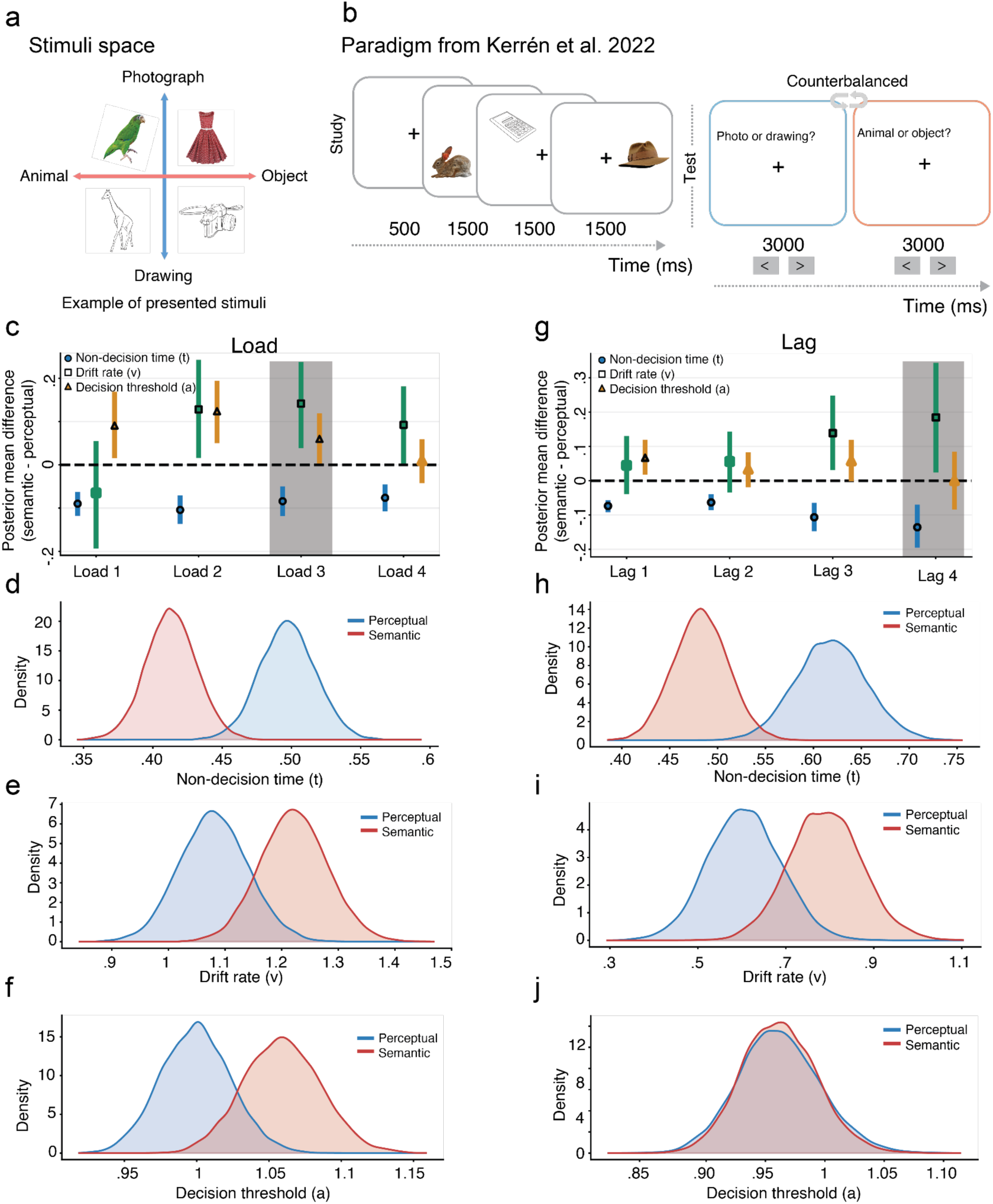
Paradigm and results of the reanalysis of Kerrén et al. **a** The stimulus space used across experiments comprised two orthogonal dimensions: a semantic dimension (animal/object) and a perceptual dimension (photograph/drawing). **b** In the paradigm from Kerrén et al. (2022), participants studied between one and four sequentially presented images and were subsequently probed on either the semantic or perceptual properties of one of the studied items. **c** Hierarchical drift-diffusion modelling revealed differences between semantic and perceptual probes in non-decision time (NDT; *t*), drift rate (*v*) and decision threshold (*a*) across memory loads. Semantic probes were associated with shorter NDT at all loads, higher drift rates at loads 2–4, and higher decision thresholds at loads 1-3. **d-f** Posterior distributions of NDT, drift rate and decision threshold, respectively, for semantic and perceptual probes at load 3 (highlighted in grey in c). Semantic probes showed shorter NDT, higher drift rates and higher decision thresholds than perceptual probes. **g** Reorganising the data according to the lag between study and test revealed shorter NDT for semantic probes at all four lags, whereas higher drift rates were supported at lags 3 and 4. A higher decision threshold for semantic probes was supported at lag 1 but not at longer lags. **h-j** Posterior distributions of NDT, drift rate and decision threshold, respectively, at lag 4 (highlighted in grey in g). Semantic probes showed shorter NDT and higher drift rates than perceptual probes, whereas decision thresholds were comparable between probe types. In c and g, points represent posterior mean semantic-minus-perceptual differences, error bars indicate 94% highest-density intervals (HDIs), and filled symbols indicate intervals excluding zero. In d-f and h-j, blue and red distributions represent perceptual and semantic probes, respectively.

Here, we reanalysed data from (Kerrén et al., 2022; Experiment 2), in which participants (n = 103 after exclusions) encoded between one and four items (the number varied pseudorandomly) at different locations on the screen. After the presentation of the items, participants were asked to recall the perceptual and semantic aspects of the stimulus they remembered in the cued location (Fig. 1B). They did so by answering yes or no to the probe questions. Load was defined by how many items they encoded in any given trial. Lag was defined as the distance between encoding and retrieval. For example, in a trial where participants saw three subsequent items at encoding, if the last item was queried during the test, that item would have lag 1, whereas the item seen first would have lag 3. We only ever analysed the first probe question when participants could not foresee which question was going to be asked.

Despite the robust behavioural effects observed in this dataset, it remains unclear what drives the semantic bias. More specifically, we hypothesised that semantic prioritisation could reflect (i) more optimal reactivation of semantic features during retrieval, (ii) selection processes operating on equally reactivated representations, or (iii) a combination of both mechanisms.

To address these questions, we applied HDD modelling to distinguish processes occurring outside evidence accumulation from those involved in decision formation. We initially considered four model specifications in which probe type (semantic versus perceptual) modulated non-decision time (*t*; Model 1), drift rate (*v*; Model 2), both parameters jointly (Model 3), or non-decision time, drift rate and decision threshold (*a*; Model 4). Non-decision time captures processes outside evidence accumulation, including stimulus encoding, preparatory processes and motor execution, whereas drift rate reflects the efficiency with which available information contributes to evidence accumulation. Decision threshold reflects the amount of evidence required before a response is made. To evaluate whether allowing decision threshold to vary improved predictive performance beyond the joint non-decision time and drift-rate model, we compared Models 3 and 4 using five-fold held-out cross-validation. At memory load 3, cross-validation favoured the full three-parameter model (Model 4; expected log predictive density [ELPD] = −2860.32) over Model 3 (ELPD = −2889.93), yielding a predictive advantage of ΔELPD = 29.61 (participant-bootstrap 95% CI [1.99, 59.84]; *P*(ΔELPD > 0) = .983). We therefore adopted the three-parameter model for the subsequent analyses, fitting it separately at each memory load and lag to characterise how the components of decision-making varied with mnemonic demands. Posterior predictive checks indicated that the model broadly reproduced the observed reaction-time distributions for semantic and perceptual judgements, including the differences between probe types (Supplementary Fig. 1A-B).

When trials were grouped by memory load, semantic judgements were associated with consistently shorter non-decision times than perceptual judgements (Fig. 1C-D). The posterior mean semantic-minus-perceptual difference was negative at every load (load 1: −90 ms, 94% highest-density interval [HDI] [−112, −68]; load 2: −104 ms, [−131, −77]; load 3: −84 ms, [−112, −55]; load 4: −76 ms, [−102, −51]), with all 94% HDIs excluding zero. Thus, the semantic advantage in non-decision time was present even when only a single item was maintained and remained evident across increasing memory loads. This suggests that semantic judgements benefit from facilitated processing outside the evidence-accumulation stage, potentially including more efficient access to task-relevant information, although non-decision time does not isolate retrieval processes specifically.

In contrast, differences in drift rate showed a more selective relationship with memory load (Fig. 1C, E). When only a single item was maintained (load 1), there was no reliable difference between semantic and perceptual judgements (mean difference = −0.065, 94% HDI [−0.188, 0.049]). At higher loads, however, semantic judgements were associated with higher drift rates than perceptual judgements (load 2: 0.128, [0.022, 0.237]; load 3: 0.142, [0.044, 0.232]; load 4: 0.092, [0.003, 0.176]), with the 94% HDI excluding zero in all three conditions. These findings suggest that the semantic advantage in non-decision time is accompanied by more efficient evidence accumulation when multiple items must be maintained concurrently. Importantly, while this pattern is consistent with a greater contribution of semantic information under increased mnemonic demands, differences between loads are descriptive, as the models were fitted separately and no formal interaction between probe type and memory load was tested.

Finally, decision thresholds also differed between semantic and perceptual judgements (Fig. 1C, F). Semantic judgements were associated with higher thresholds at loads 1-3 (load 1: 0.091, 94% HDI [0.021, 0.163]; load 2: 0.124, [0.056, 0.189]; load 3: 0.060, [0.008, 0.113]), whereas the difference at load 4 was negligible (0.010, [−0.037, 0.054]). Higher decision thresholds for semantic judgements indicate that participants required more accumulated evidence before responding, potentially counteracting the response-time benefits associated with shorter non-decision times or higher drift rates. Together, these results suggest that semantic prioritisation is not attributable to a single decision component. Rather, semantic judgements show a consistent advantage in non-decision time across memory loads, alongside differences in evidence accumulation and decision thresholds that vary with the number of maintained items.

When trials were grouped by the lag between study and test, we examined whether the semantic advantage in non-decision time persisted across lags and whether differences in evidence accumulation emerged primarily at longer delays. We additionally examined decision thresholds to determine whether semantic and perceptual judgements differed in the amount of evidence required before a response was made. Consistent with the preceding analyses, semantic judgements were associated with shorter non-decision times than perceptual judgements at all four lags (Fig. 1G, H). The posterior mean difference (semantic minus perceptual) was negative at every lag (lag 1: −74 ms, 94% highest-density interval [HDI] [−92, −57]; lag 2: −63 ms, [−86, −39]; lag 3: −107 ms, [−147, −65]; lag 4: −136 ms, [−195, −70]), with all 94% HDIs excluding zero. These findings suggest a consistent semantic advantage in processes captured by non-decision time across study-test delays, with the numerically largest difference observed at the longest lag.

In contrast, drift-rate differences showed a more selective pattern across lags (Fig. 1G, I). At shorter lags, differences between semantic and perceptual judgements were small and uncertain (lag 1: mean difference = 0.045, 94% HDI [−0.039, 0.130]; lag 2: 0.056, [−0.035, 0.143]), with both intervals overlapping zero. At longer lags, however, semantic judgements were associated with higher drift rates than perceptual judgements (lag 3: 0.139, 94% HDI [0.031, 0.248]; lag 4: 0.184, [0.024, 0.344]), with both intervals excluding zero. Thus, although semantic judgements were associated with shorter non-decision times across all lags, an additional advantage in evidence accumulation was supported at longer study-test delays.

Finally, we examined decision thresholds to determine whether semantic and perceptual judgements differed in the amount of evidence required to reach a decision (Fig. 1G, J). At lag 1, semantic judgements were associated with a higher decision threshold than perceptual judgements (mean difference = 0.067, 94% HDI [0.018, 0.119]). At subsequent lags, however, the differences were less conclusive (lag 2: 0.033, [−0.019, 0.083]; lag 3: 0.057, [−0.002, 0.119]; lag 4: −0.000, [−0.084, 0.084]), with all three intervals overlapping zero. Thus, whereas semantic judgements required more evidence before responding at the shortest lag, there was no clear evidence for a corresponding difference in decision thresholds at longer delays.

Together, these results indicate that semantic prioritisation in working memory is expressed through distinct components of the decision process. Shorter non-decision times for semantic relative to perceptual judgements were consistently observed across all memory loads and study-test lags, indicating an advantage in processes occurring outside evidence accumulation. In contrast, higher drift rates for semantic judgements were supported at loads 2-4 and at longer study-test lags (3-4), suggesting an additional advantage in the efficiency of evidence accumulation when multiple items must be maintained or information must be retrieved after longer delays. Decision thresholds were also higher for semantic judgements at loads 1-3 and at the shortest lag, but these differences were not consistently observed across conditions. This dissociation suggests that semantic prioritisation may primarily reflect facilitated access to stored representations or other processes preceding evidence accumulation, with additional contributions from accumulation efficiency under greater mnemonic demands. However, because non-decision time also encompasses processes unrelated to retrieval, and models were fitted separately across loads and lags, these interpretations remain tentative and differences between conditions should be considered descriptive rather than evidence of formal interactions.

### Experiment 1

The drift-diffusion modelling results obtained from re-analysing our previous dataset suggest that semantic prioritisation in working memory arises from at least two separable components: (i) an advantage in processes occurring before evidence accumulation for semantic information, reflected in shorter non-decision times across loads and lags, and (ii) a more selective advantage in evidence accumulation that emerges under higher cognitive demands. However, these analyses alone cannot determine why semantic information is prioritised. In particular, it remains unclear whether the observed semantic advantage reflects a genuine retrieval-related benefit, where semantic representations are more readily reactivated from working memory, or whether it instead arises from a selection bias operating after information has already been accessed.

Experiment 1 was designed and pre-registered (https://researchbox.org/5966) to adjudicate between these possibilities by manipulating the extent to which retrieval demands were required at the time of the decision (Fig. 2A). By introducing a retro-cue that either specified the relevant memory item in advance or remained neutral, we aimed to dissociate retrieval- related processes from selection at the decision stage (Griffin & Nobre, 2003; Kuo et al., 2012). In trials where a valid retro-cue was presented prior to the probe, we hypothesised that item selection/prioritisation or reactivation could commence prior to the probe. In contrast, in neutral trials where the retro-cue was uninformative, retrieval and selection demands must both be resolved after the probe, such that any observed semantic advantage could reflect differences in either or both processes. If semantic prioritisation primarily reflects differences in retrieval, then reducing selection uncertainty and allowing advance prioritisation of the relevant item via a valid cue should attenuate or eliminate the semantic advantage. Conversely, if semantic prioritisation reflects selection biases operating on already retrieved information, then the advantage should persist even when retrieval demands are minimised. In the following experiment, we combined behavioural measures with drift-diffusion modelling to examine how semantic and perceptual information are differentially accessed and accumulated under valid and neutral cue-conditions.

**Figure 2.**
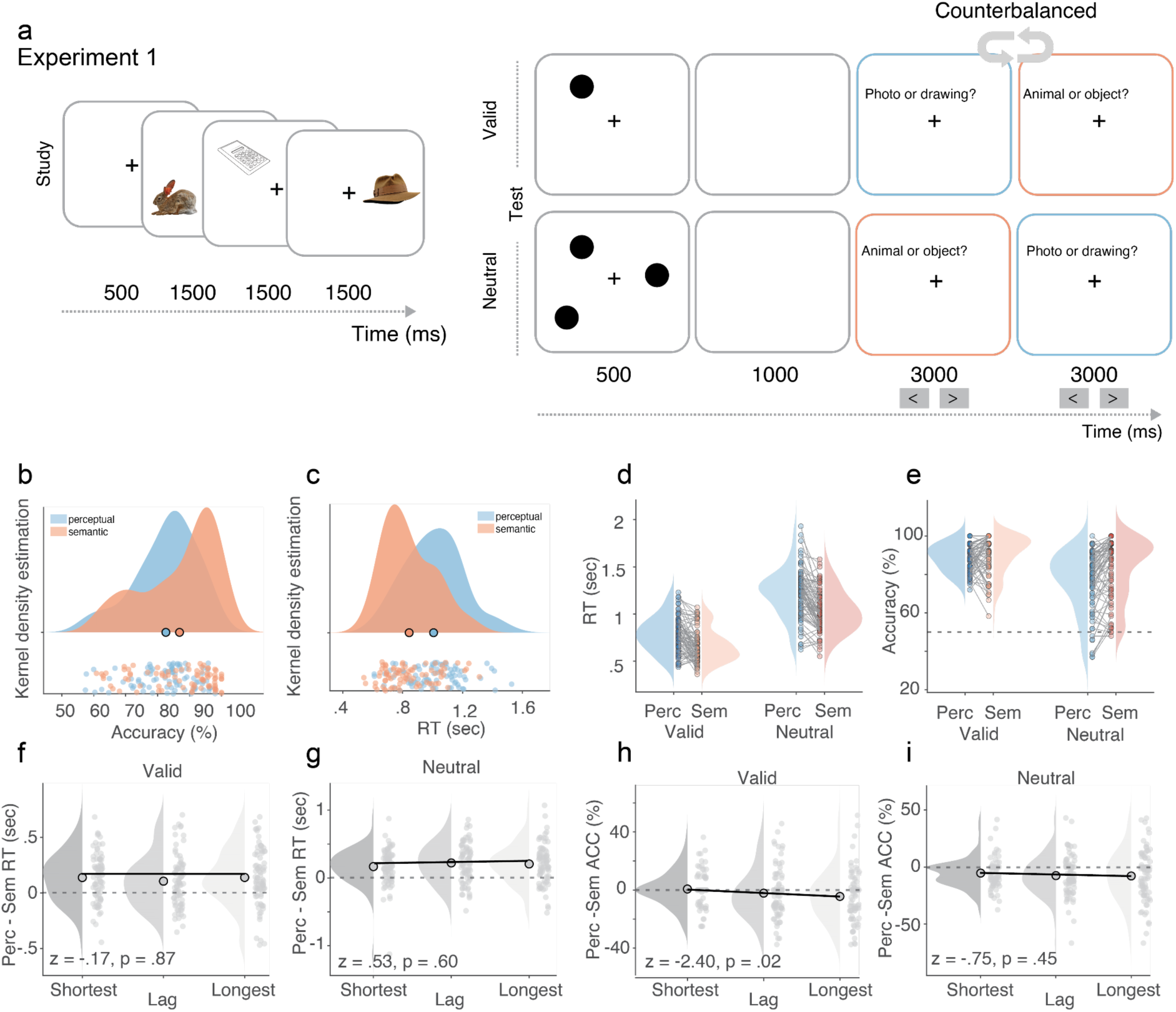
Paradigm and results of Experiment 1. **a** Behavioural paradigm. During study, participants saw three items sequentially in pseudorandom positions around the fixation cross. After encoding, they either saw a single circle (valid condition), indicating which item was going to be tested later, or three circles (neutral condition), indicating the three positions in which items were presented during study. Following a 1000ms blank screen, they were tested for the cued item (in the valid condition), or a random item (neutral condition). **b** Average accuracy for semantic trials was higher than perceptual trials across conditions. **c** Average reaction times (RTs) for semantic trials were faster than perceptual trials. **d** We observed faster semantic RTs in both valid and neutral conditions. **e** We observed higher accuracy for the semantic questions in the neutral condition, but not in the valid one. **f** No difference between conditions for valid trials. **g** And not for neutral trials either. **h** A significant negative slope was found for the valid condition, indicating that semantic prioritisation increased with lag. **i** This was not the case in neutral trials.

Eighty-one participants were exposed to 3 items during study, sequentially presented in three pseudorandom locations on the screen (each for 1500ms). We selected three items because this condition yielded the most reliable differences in non-decision time, drift rate and decision threshold in the reanalysis of our previous dataset. The task for participants was to remember the items and the locations. Immediately after the third item, a retro-cue was shown. In the valid condition, a circle was shown in the location where the questions about the item was going to be shown. In the neutral condition, three circles were shown in the location of the three studied items. Following a 1000ms delay, a question appeared on screen (“Animal or Object?” or “Photo or Drawing?”, counterbalanced; see Methods) for three seconds. Participants were instructed to push the right or left button to register a response. Once pushed, the font colour of the question changed to either green (correct) or red (incorrect). Thereafter the next question appeared. If the first question probed the semantic dimension, e.g., “Animal or Object?”, the second question probed the perceptual features, e.g., “Photo or Drawing?” (counter-balanced; see Methods). We analysed accuracy and reaction times (RTs) to the first question of each trial.

Participants were on average good at performing the task (accuracy: semantic question mean (M) = 86.85%, standard deviation (SD) = 11.64%; perceptual question M = 82.58%, SD = 9.73%) and were on average better in the semantic as compared to the perceptual question (t(1,80) = 4.71, p < .01, Bayesian t-test against zero comparing semantic and perceptual probes: BF10 > 1000; Fig. 2B). They were also faster at responding to the semantic as compared to the perceptual questions (semantic: M = 844ms, SD = 172ms; perceptual: M = 1007ms, SD = 191ms; t(1,80) = −9.85, p < .01, BF10 > 1000; Fig. 2C). These results very much mimic previous results (Kerrén et al., 2022) and suggest that categorical semantic information is overall easier and faster to remember.

Our main goal of Exp. 1 was to understand how much the semantic prioritisation was due to a selection bias as compared to activation in advance of decision processes. In the valid condition, the retro-cue signalled which of the three items was going to be probed, thereby allowing participants to pre-activate it. In the neutral condition, such pre-activation was not possible. Instead, participants had to keep all three items simultaneously in mind. Our main hypothesis was that there would be more semantic prioritisation in the neutral condition, where memory retrieval is required during the probe, as compared to the valid condition. Indeed, analysing RTs using a Bayesian t-test against zero, we observed moderate evidence (BF10 = 5.31; Fig. 2D) for more semantic prioritisation (larger difference between RTs for semantic and perceptual question) for the neutral as compared to the valid condition. Although our results suggest that selection had an impact on the format of working memory representations, we still found a reliable semantic prioritisation in both conditions, separately (valid: mean difference semantic and perceptual RTs = 128ms, SD = 171ms, BF10 > 1000; neutral: mean difference semantic and perceptual RTs = 198ms, SD = 201ms, BF10 > 1000), indicating that memory reactivation of previously studied items adhere to earlier availability of the semantic features.

Also for accuracy did we observe that semantic features were more favoured in the neutral condition (difference semantic and perceptual accuracy in valid condition M = 1.75%, SD = 10%; neutral condition M = 6.76%, SD = 10.43%, BF10 = 68.36; Fig. 2E). Interestingly, when analysing each condition separately, we did not find any evidence for a semantic prioritisation in the valid condition (semantic M = 90.5%, SD = 10%; perceptual M = 88.75%, SD = 6.92%, BF10 = .41). In contrast, we did observe it in the neutral condition (semantic M = 83.28%, SD = 15.70%; perceptual M = 76.51%, SD = 15.61%, BF10 > 1000).

To further understand the semantic prioritisation, we split the data into lag between study and test. As the items were presented sequentially during study, the lastly presented item during study would be regarded the closest in time to test, whereas the firstly presented item during study would be the one farthest away. In our previous study we found that when there was short time between study and test, we observed less semantic prioritisation as compared to when the lag was larger (Kerrén et al., 2022). We did not find any evidence for a positive slope in any of the conditions when assessing RTs (valid difference distance 1: M = 138ms, SD = 215ms; distance 2: M = 105ms, SD = 231ms; distance 3: 139ms, SD = 249ms, z = −.17, p = .87; neutral difference distance 1: M = 168ms, SD = 261ms; distance 2: M = 222ms, SD = 281ms; distance 3: 206ms, SD = 287ms, z = .53, p = .60; Fig. 2F and 2G). However, when assessing accuracy, we did find a negative slope for the valid condition (valid difference distance 1: M = .71%, SD = 11.69%; distance 2: M = −2.17%, SD = 16.98%; distance 3: − 4.41%, SD = 17.18%, z = −2.40, p = .02; neutral difference distance 1: M = −5.31, SD = 12.83%; distance 2: M = −7.53%, SD = 17.30%; distance 3: −7.84%, SD = 21.89%, z = −.75, p = .45; Fig. 2H and 2I). Interestingly, we found numerically higher accuracy for perceptual probes when study and test were closest in time in the valid condition, however, with moderate evidence for the null-hypothesis (BF10 = .14). Together, these results suggest that lag-related modulation of semantic prioritisation depends on the ability to focus attention on a single item, with time effects emerging only under valid cueing and not when attention must remain distributed across items.

#### Balanced integration score

To take into account the speed-accuracy trade-off, we used the balanced integration score (BIS), which quantifies performance in terms of both RT and accuracy with equal weight (Liesefeld et al., 2015). Here we calculated the difference in the BIS between perceptual and semantic probes where larger values reflect more semantic prioritisation. As predicted, we found strong evidence for the more semantic prioritisation in neutral condition compared to the valid condition (valid: M = .97, SD = 1.46; neutral: M = 1.91, SD = 1.88, both BF10 > 1000 when tested against zero and BF10 = 984 when tested for difference between valid and neutral; Fig. 3a). For valid trials, this result was likely driven by the faster RTs to semantic as compared to the perceptual probes.

**Figure 3.**
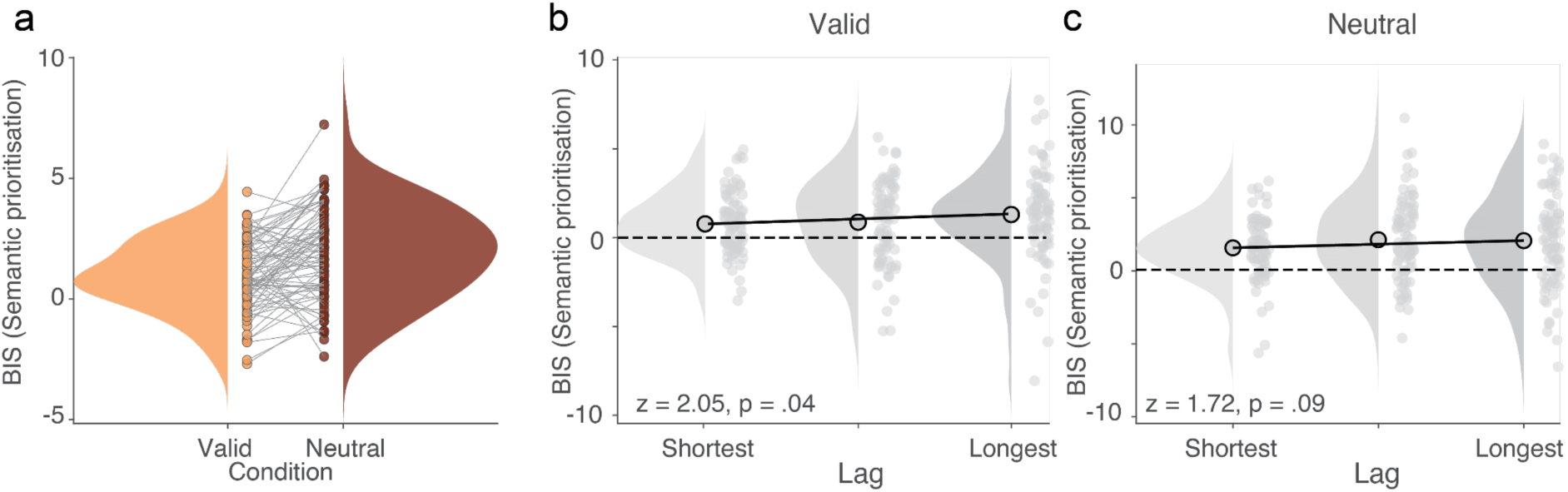
Balanced integration score. **a** BIS was significantly higher in neutral trials (brown) as compared to valid trials (orange). **b** When organising the data in the valid condition according to lag between study and test, we observed a significant positive slope across lags, reflecting more semantic prioritisation for larger lags. **c** This was not the case for the neutral condition, where a similar semantic prioritisation was present in all lag levels.

When evaluating distances, we found a significant positive slope for the valid condition (z = 2.05, p = .04; Fig. 3B), suggesting that when participants could focus on one item, the semantic prioritisation was reduced, but then increased with lag. However, separate tests still revealed significant semantic prioritisation for each distance (distance 1: M = .78, SD = 1.71, t(1,80) = 4.11, p < .01; BF10 = 208; distance 2: M = .87, SD = 2.28, t(1,80) = 3.46, p < .01; BF10 = 27.21; distance 3: M = 1.30, SD = 2.54, t(1,80) = 4.65, p < .01; BF10 > 1000). We did not find a significant slope for the neutral condition (z = 1.72, p = .09; ; Fig. 3C), suggesting a similar semantic prioritisation for all distance levels (distance 1: M = 1.57, SD = 2.10, t(1,80) = 6.79, p < .01; BF10 > 1000; distance 2: M = 2.13, SD = 2.77, t(1,80) = 6.96, p < .01; BF10 > 1000; distance 3: M = 2.07, SD = 3.13, t(1,80) = 5.99, p < .01; BF10 > 1000).

Taken together, Exp.1 shows that the observed semantic prioritisation in VMW may be related to a memory reactivation effect. However, allowing participants to pre-activate one of the three items from study has a significant effect, but it does not eliminate the semantic prioritisation completely.

#### Drift-diffusion modelling of Experiment 1

Together, the behavioural results of Experiment 1 indicate that semantic prioritisation in working memory is robust, but sensitive to attentional demands. Valid retro-cues reduced the magnitude of the semantic advantage, consistent with a contribution from processes involved in selecting and accessing relevant memory representations. Nevertheless, semantic judgements remained faster than perceptual judgements in both cueing conditions, suggesting that semantic prioritisation cannot be explained exclusively by differences in retrieval demands. Instead, the behavioural advantage may reflect differences in how semantic and perceptual information contribute to multiple stages of decision formation.

Reaction time and accuracy alone cannot establish whether these differences arise from processes preceding evidence accumulation, from the efficiency of evidence accumulation itself, or from differences in decision strategy. To characterise these contributions, we applied hierarchical drift-diffusion modelling, allowing feature type (semantic vs perceptual) and cue validity (valid vs neutral) to modulate non-decision time (*t*), drift rate (*v*), and decision threshold (*a*). We compared models in which these parameters varied individually or jointly (see Supplementary Fig. 1C, D for posterior predictive checks). Similar to the reanalysis of our previous experiment, also in Exp. 1 five-fold cross-validation favoured the full model, allowing all three parameters to vary, over the simpler model incorporating only non-decision time and drift rate (held-out ELPD difference = 107.92, participant-bootstrap 95% CI [64.06, 155.44]). We therefore report results from the full model.

If semantic prioritisation reflects an advantage in processes preceding evidence accumulation that is influenced by retrieval demands, we would expect a larger semantic-perceptual difference in non-decision time under neutral cueing, where the relevant memory item cannot be prioritised in advance, than under valid cueing, where item selection and reactivation can begin before the probe. Conversely, if semantic prioritisation also involves differences in the efficiency with which information contributes to evidence accumulation, we would expect higher drift rates for semantic judgements. Whether this accumulation advantage is itself modulated by cue validity provides a further test of the extent to which attentional prioritisation influences distinct components of decision formation.

For non-decision time, we observed a pronounced feature-by-validity interaction (Fig. 4A, D). Under neutral cueing, semantic judgements were associated with substantially shorter non- decision times than perceptual judgements (semantic minus perceptual = −.118 s, 94% highest density interval [−.142, −.096], posterior probability < 0 = 1). Under valid cueing, this advantage was largely eliminated (−.003 s, 94% highest density interval [−.014, .008], posterior probability < 0 = .701). Correspondingly, the interaction was positive and credibly different from zero (change from neutral to valid = .115 s, 94% highest density interval [.091, .141], posterior probability > 0 = 1). These findings indicate that the semantic advantage in processes captured by non-decision time is strongly attenuated when attentional prioritisation is possible, consistent with facilitated access to semantic information when the relevant representation must be selected or reactivated at the time of the probe.

**Figure 4.**
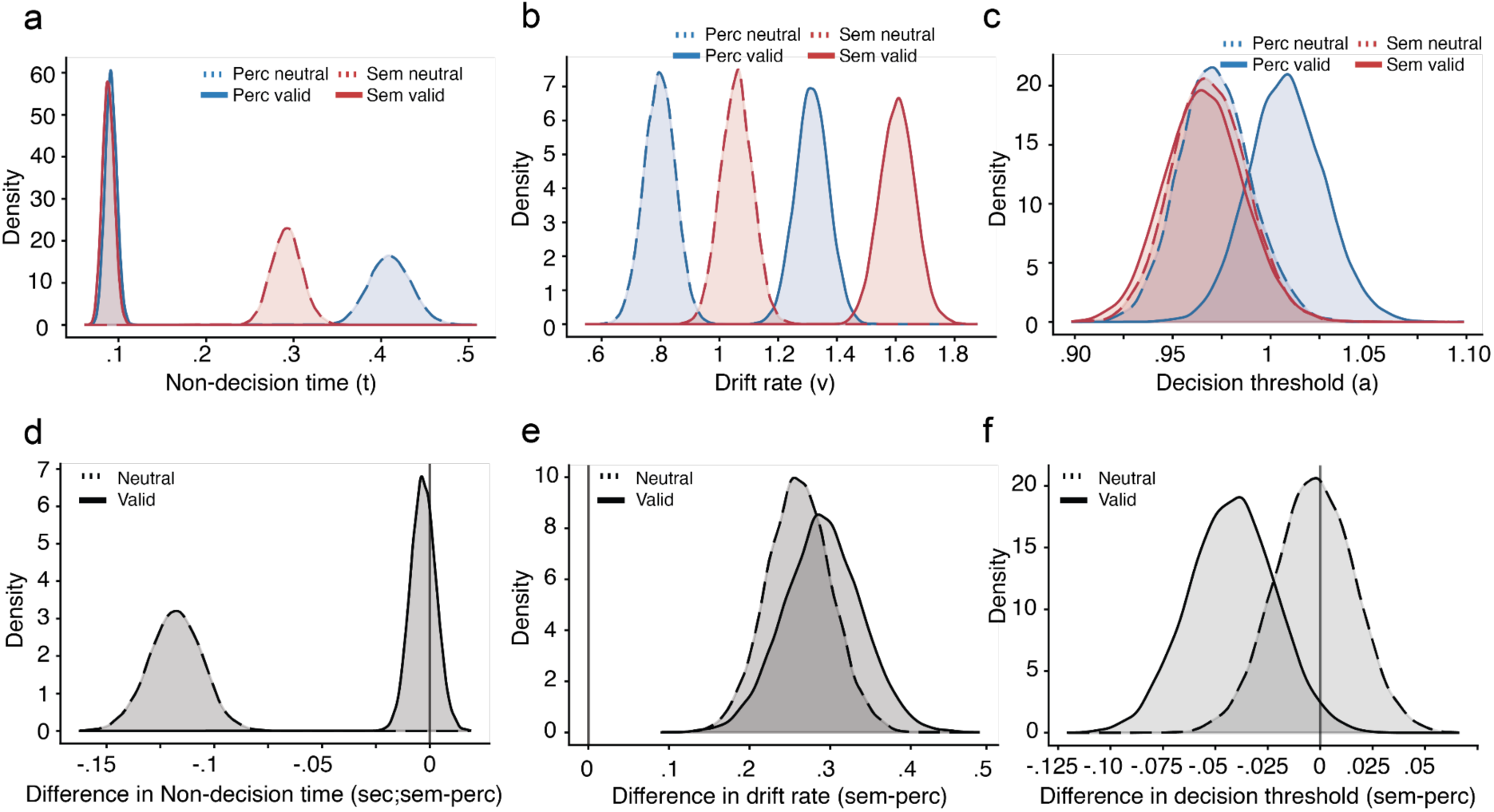
Drift-diffusion modelling of Experiment 1. Posterior distributions from the hierarchical drift- diffusion model allowing non-decision time, drift rate and decision threshold to vary as a function of feature type and cue validity. **a** Non-decision time was substantially shorter for semantic (red) than perceptual (blue) judgements in the neutral condition, whereas this difference was largely eliminated under valid cueing. **b** Drift rates were higher for semantic than perceptual judgements in both cueing conditions, with no reliable interaction between feature type and cue validity. **c** Decision thresholds were similar for semantic and perceptual judgements under neutral cueing, whereas semantic judgements showed a small reduction in threshold under valid cueing. **d-f** Corresponding posterior distributions of the semantic-minus-perceptual differences in non-decision time, drift rate and decision threshold, respectively. Throughout, solid lines represent valid trials and dashed lines represent neutral trials. The vertical line at zero in the bottom panels indicates no difference between feature types.

For drift rate, semantic judgements were associated with higher rates of evidence accumulation in both cueing conditions (Fig. 4B, E). Under neutral cueing, semantic drift rates were reliably higher than perceptual drift rates (semantic minus perceptual = .261, 94% highest density interval [.182, .334], posterior probability > 0 = 1). A comparable advantage was observed under valid cueing (.291, 94% highest density interval [.200, .383], posterior probability > 0 = 1). However, in contrast to non-decision time, the feature-by-validity interaction was small and uncertain (change from neutral to valid = .029, 94% highest density interval [−.087, .144], posterior probability > 0 = .683). Thus, while semantic information contributed more efficiently to evidence accumulation across both conditions, we found no reliable evidence that this advantage was selectively enhanced by valid cueing.

Allowing decision threshold to vary did not eliminate the semantic advantage in non-decision time (Fig. 4C, F). Under neutral cueing, semantic and perceptual judgements showed similar thresholds (semantic minus perceptual = −.003, 94% highest density interval [−.040, .031]). Under valid cueing, semantic judgements were associated with a small reduction in threshold (−.042, 94% highest density interval [−.082, −.005], posterior probability < 0 = .980). However, the feature-by-validity interaction remained uncertain (−.039, 94% highest density interval [−.091, .013]), providing no clear evidence that cue validity systematically modulated the relative decision thresholds for semantic and perceptual information.

Together, these results suggest that semantic prioritisation in working memory involves dissociable contributions from processes outside evidence accumulation itself. Semantic judgements were associated with higher drift rates irrespective of cue validity, whereas their pronounced advantage in non-decision time emerged primarily when the relevant memory representation could not be prioritised in advance. Importantly, this effect persisted when differences in decision threshold were accommodated, suggesting that it cannot be explained solely by differences in decision caution. Although non-decision time does not uniquely identify memory retrieval, the selective attenuation of the semantic advantage under valid cueing is consistent with more efficient access to semantic than perceptual information when stored representations must be selected or reactivated from an unattended state.

### Experiment 2

Results from Experiment 1 showed that attentional prioritisation differentially influences the decision processes underlying semantic and perceptual judgements. Valid retro-cueing substantially attenuated the semantic advantage in non-decision time, whereas higher drift rates for semantic judgements remained evident in both cueing conditions. Thus, the availability of information about the relevant memory item beforehand primarily altered the semantic-perceptual difference in processes outside evidence accumulation, without reliably changing the relative efficiency of evidence accumulation. A remaining question is whether these effects depend on attentional interference during maintenance or on the passage of time itself.

A remaining question is whether semantic prioritisation depends on the need to actively maintain and protect memories over time or from interference (Oberauer & Lin, 2017). Experiment 2 was designed and pre-registered (https://researchbox.org/5966) to address this issue by manipulating both the temporal delay between study and test and the presence of intervening interference, allowing us to examine whether semantic prioritisation is selectively enhanced when visual working memory content must be shielded from competing cognitive demands.

To do so, we designed a task in which we varied time between study and test, and interference of a competing task (see: Padezhki et al., (2025) and van Loon et al., (2018) for a similar manipulation). A total of 77 participants took part in the experiment which had three conditions. The *Immediate* condition mirrored the neutral condition of Exp. 1, such that after having studied three items, participants were shown a screen saying “Now”, signalling that retrieval of one of the three items would start immediately. After a short delay (1000ms), the same questions as in Exp. 1 appeared subsequently. In the *Prospective* condition, participants were shown a screen with the word “Later”, whereafter a short delay of 1000ms, they saw a symbol of a triangle, circle or square, together with the word triangle?, circle?, or square? and were asked to respond whether the symbol matched the word (2000ms). After this intervening task, they answered the questions related to one of the items from the study phase. In the *Delayed* condition, participants instead were shown a screen with the word “Wait”, whereafter a delay task (1000ms) they were shown a merged symbol of the triangle, circle and square, and were asked to do nothing for the 2000ms it was shown. After this they answered the questions regarding one of the studied items (Fig. 5A).

**Figure 5.**
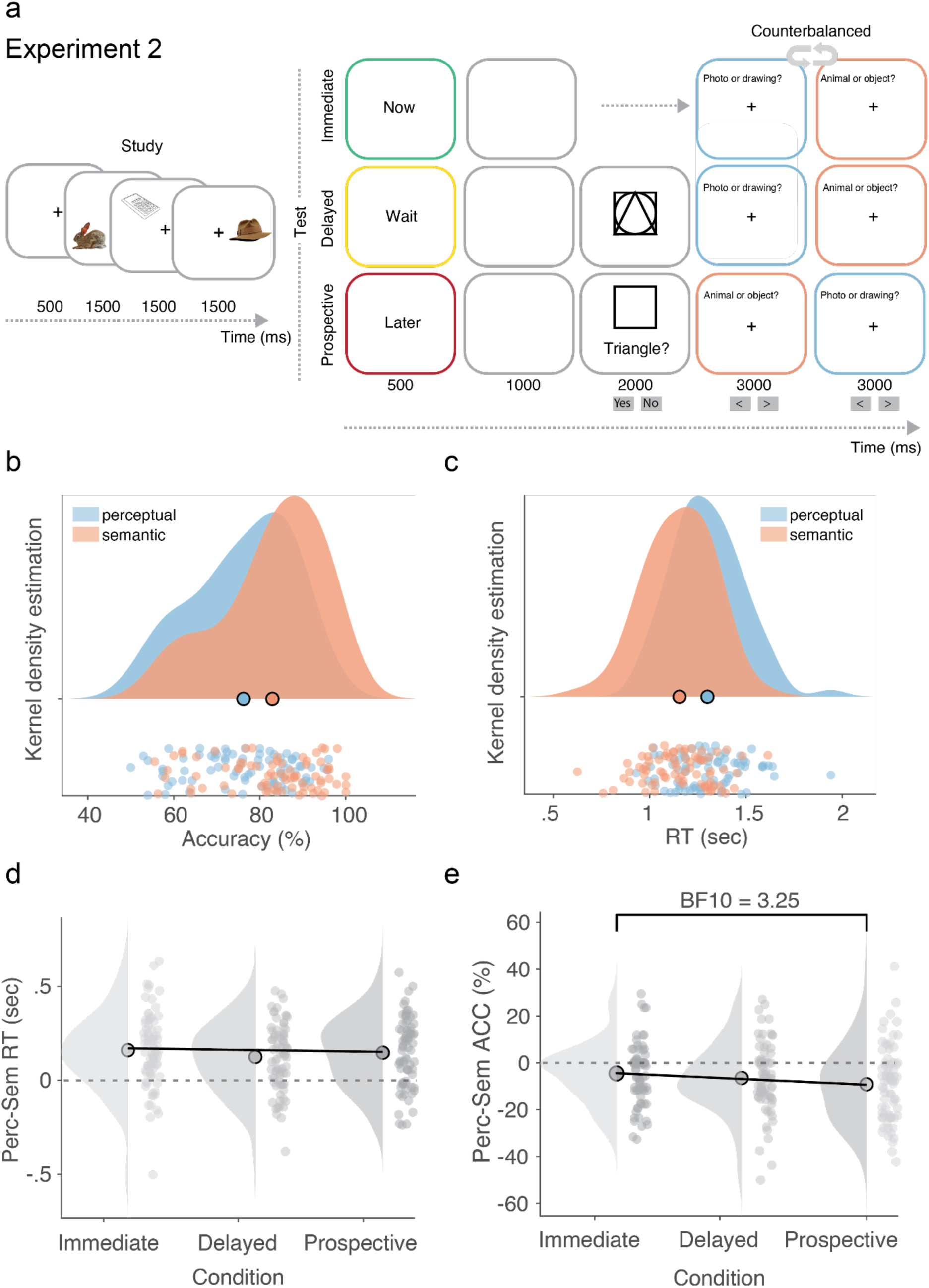
Paradigm and results of Experiment 2. **a** The study phase was identical to Exp. 1. After encoding, participants saw three different words (“Now”, “Wait”, or “Later”), indicating which condition they were in (Immediate, Delayed, or Prospective). In the Immediate condition, they were randomly tested on one of the presented items from the study phase, immediately following a brief 1000ms blank screen. In the Delayed condition, participants were exposed to a symbol consisting of a triangle, square and a circle, stacked on top of each other for 2000ms. After this, they were randomly questioned about one of the studied items. In the Prospective condition, participants saw a triangle, square or a circle with the question “Triangle?”, “Square?”, or “Circle?”, and were asked to respond yes if the word matched the symbol, and no otherwise. After 2000ms, the trial continued in the same manner as the two other conditions. In the Immediate condition, we asked participants the same symbol matching question after they had responded to the questions about the item from encoding (not shown in figure). **b** Average accuracy for semantic trials was higher than perceptual trials. **c** Average reaction times (RTs) for semantic trials were faster than perceptual trials. **d** We observed faster semantic RTs in all conditions, but no difference between conditions. **e** The difference between semantic and perceptual accuracy was lowest for the Immediate condition, followed by Delayed and Prospective, with a significant difference between Immediate and Prospective conditions.

Critically, the three conditions were designed to dissociate the effects of temporal delay from the need to actively shield working memory representations from interference. Comparing the *Immediate* and *Delayed* conditions allowed us to examine the consequences of extending the retention interval without introducing an additional response requirement before the memory probe. Comparing the *Delayed* and *Prospective* conditions assessed the additional effects of performing an intervening task during maintenance, while holding the overall retention interval constant. Although this comparison was designed to manipulate interference with working- memory maintenance, it also introduced decision-making, response preparation and task- switching demands. These factors therefore cannot be fully dissociated within the present design.

Participants were on average performing better in the semantic question (M = 82.92%, SD = 11.77%) as compared to the perceptual question (M = 76.22%, SD = 11.71%, t(1,76) = 5.95, p < .01, BF10 > 1000; Fig 5B). They were also faster at responding to the semantic as compared to the perceptual questions (semantic: M = 1150ms, SD = 180ms; perceptual: M = 1300ms, SD = 184ms; t(1,76) = 9.49, p < .01, BF10 > 1000; Fig 5C). Again, semantic features are consistently easier and faster to remember. Moreover, the new experimental conditions introduced in this experiment significantly impacted behaviour (main effect of condition in accuracy: F = 73, p < .001, BF > 1000; and in RT: F = 223, p < .001, BF > 1000). Participants were most accurate in the Immediate condition (M = 85.34%, SD = 13.4%) compared to the Delayed (M = 81.08%, SD = 13.66%, BF10 > 134) and the Prospective ones (M = 72.28%, SD = 14.86%, BF10 > 1000). Interference also impacted behaviour above and beyond the passing of time, as revealed by the greater accuracy in Delayed compared to Prospective (BF10 > 1000). A similar pattern was observed in RT, with fastest responses in Immediate (M = 1069ms, SD = 199ms) compared to Delayed (M = 1128ms, SD = 201ms, BF > 1000) and Prospective (M = 1396ms, SD = 227ms, BF > 1000). Prospective yielded slower responses compared to Delayed trials as well (BF > 1000).

Our main hypothesis was that if semantic prioritisation is driven by shielding against interfering processes, we would see an increase in semantic prioritisation in the *Prospective* condition as compared to the *Immediate* and *Delayed* condition. Our second hypothesis regarded time between study and test. In our previous paper (Kerrén et al., 2022), we observed an increase in semantic prioritisation the further away the tested item was from study. However, this measure was confounded by varying the number of items presented at study. Therefore, in the present task, we kept the number of items studied to 3. If time further transformed the memory trace, we would see an increase in semantic prioritisation in the delayed as compared to the immediate condition.

To test these hypotheses, we conducted a Bayesian repeated-measures ANOVA with Condition (Immediate, Prospective, Delayed) as a within-subject factor, applied separately to reaction time (RT) and accuracy. Bayesian model comparison provided moderate evidence for the null model, indicating no effect of Condition on RT (BF = .12). Nevertheless, since we had a pre-registered directional hypothesis, we directly tested *Immediate* vs. *Prospective* and, contrary to our prediction, found strong evidence for the null hypothesis (Bayesian t-test against zero comparing Immediate and Prospective = .09, paired-samples t-test: t(1,76) = −.51, p = .70, one-sided; difference semantic and perceptual question Immediate condition: M = 160ms, SD = 198ms; Prospective condition: M = 146ms, SD = 182ms; Fig. 5D).

An identical repeated-measures ANOVA, but now using accuracy, showed inconclusive evidence (BF = .46). Again, we directly tested the pre-registered directional hypothesis of a difference between *Immediate* and *Prospective* condition. In this case we found moderate evidence for more semantic prioritisation in the prospective as compared to the immediate condition (Bayesian t-test against zero comparing Immediate and Prospective = 3.25; t(1,76) = 2.35, p = .01, one-sided; difference semantic and perceptual question Immediate condition: M = −4.5%, SD = 12.61%; Prospective condition: M = −9.13%, SD = 15.77%; Fig. 5E), suggesting that a larger delay plus interference from another task increases the semantic prioritisation, reflected in the accuracy of the retrieved memory.

Our second directional hypothesis was that the *Delayed* condition should show more semantic prioritisation as compared to the *Immediate* condition if time between study and test has an effect on the format of working-memory representations. A comparison between *Delayed* and *Prospective* conditions would show an additional effect of task interference. On the contrary to this prediction, we observed numerically more semantic prioritisation in the *Immediate* (M = 160ms, SD = 198ms) as compared to the *Delayed* (M = 125ms, SD = 176ms) condition when assessing RTs, with the Bayesian t-test against zero showing strong evidence for the null- hypothesis (BF = .05; t(1.76) = −1.46, p = .93, one-sided). When contrasting *Delayed* and *Prospective* conditions, we found strong evidence for the null hypothesis (BF10 = .07), such that participants had numerically more semantic prioritisation in the *Prospective* condition (M = 147ms, SD = 182ms). For accuracy, we found numerically higher semantic prioritisation in the *Delayed* (M = −6.46%, SD = 14.50%) as compared to the *Immediate* (M = −4.50%, SD = 12.61%). However, Bayesian t-test against zero comparing *Immediate* and *Delayed* showed moderate evidence for the null hypothesis (BF = .32, t(1,76) = .96, p = .17, one-sided). When contrasting *Delayed* and *Prospective* conditions, we again found numerically more semantic prioritisation in the Prospective condition (M = −9.13%, SD = 15.77%), yielding strong evidence for the null hypothesis (BF10 = .06).

#### Balanced integration score

We again used BIS to understand the speed-accuracy trade-off. Immediate (M = 1.57, SD = 2.03) and Delayed (M = 1.50, SD = 2.01) conditions had numerically less semantic prioritisation than the Prospective condition (M = 1.92, SD = 2.12). Using a one-way Bayesian ANOVA, we found strong evidence for a null effect (BF10 = .015). Our directional follow-up test testing for a difference in BIS between Immediate and Prospective conditions revealed inconclusive evidence (BF = .50, t(1,76) = 1.29, p = .1, one-sided), whereas the test between Immediate and Delayed conditions showed moderate to strong evidence for the null hypothesis (BF = .10, t(1,76) = −.26, p = .60, one-sided).

Across behavioural measures, the results provided limited support for the hypothesis that semantic prioritisation in visual working memory is selectively enhanced by temporal delay alone, but partial support for a role of interference- and delayed-related demands. Reaction time measures showed consistent evidence for the absence of condition effects, with semantic prioritisation in RTs remaining stable across *Immediate*, *Delayed*, and *Prospective* conditions. In contrast, accuracy revealed moderate evidence for increased semantic prioritisation in the *Prospective* condition relative to the *Immediate* condition, suggesting that the presence of an intervening task, requiring the maintenance and protection of memory representations, can enhance the semantic advantage. No reliable differences were observed between *Immediate* and *Delayed* conditions in either RT or accuracy, providing evidence against a simple effect of time between study and test when memory load is held constant. Finally, analyses of the balanced integration score yielded evidence consistent with the null hypothesis, indicating that the observed effects were not driven by systematic speed-accuracy trade-offs. Together, these findings suggest that semantic prioritisation is not uniformly amplified by delay, but may be selectively strengthened when working memory representations must be actively shielded from interference compared to an immediate implementation.

#### Drift-diffusion modelling of Experiment 2

Together, these findings suggest that interference and time may differentially affect the components of working memory performance, but that these effects are not readily captured by aggregate behavioural measures alone. To characterise the decision processes underlying semantic prioritisation in Experiment 2, we fitted a hierarchical drift-diffusion model in which feature type (semantic versus perceptual), condition (Immediate, Prospective, Delayed) and their interaction jointly modulated non-decision time (*t*), drift rate (*v*) and decision threshold (*a*). Based on the superior predictive performance of the three-parameter model in Experiment 1 and in the reanalysis of Kerrén et al. (2022), we adopted this model for Experiment 2. This allowed us to examine whether semantic prioritisation was expressed through differences in processes occurring outside evidence accumulation, the efficiency of evidence accumulation itself, or the amount of evidence required before responding. Posterior predictive checks further indicated that the model broadly reproduced the observed reaction-time distributions across experimental conditions and probe types (Supplementary Fig. 1E-G).

Across all conditions, semantic judgements were associated with reliably shorter non-decision times than perceptual judgements, indicating an advantage in processes occurring outside evidence accumulation (Fig. 6a). This semantic advantage was present in the *Immediate* condition (mean difference in *t* = −86 ms, 94% highest-density interval [HDI] [−108, −64]), the *Prospective* condition (−134 ms, [−164, −104]) and the *Delayed* condition (−94 ms, [−122, −69]). In all three conditions, the posterior probability that the semantic–perceptual difference was negative approached 1, providing strong evidence for a consistent semantic advantage in non- decision time.

**Figure 6.**
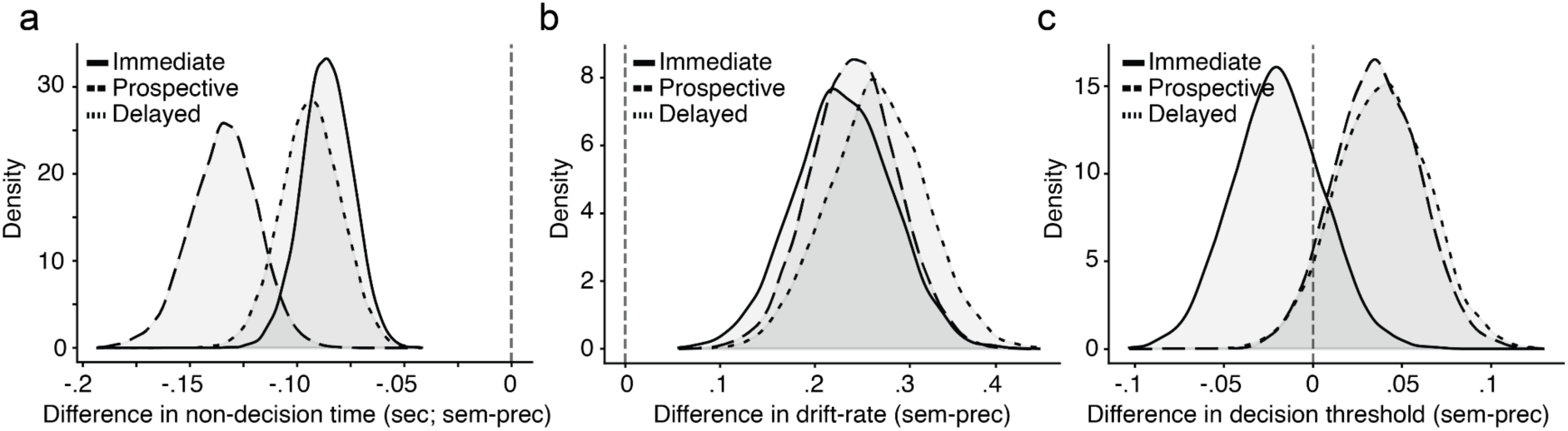
Drift diffusion modelling of Experiment 2. Semantic judgements were associated with shorter non-decision times across all conditions, with a larger absolute advantage in the Prospective condition compared with Immediate and Delayed. **b** Semantic judgements showed higher drift rates across all three conditions, but the magnitude of this advantage did not reliably differ between conditions. **c** Decision thresholds did not reliably differ between semantic and perceptual judgements in any condition. Solid, dashed and dotted lines represent Immediate, Prospective and Delayed conditions, respectively. Vertical dashed lines indicate zero difference.

Importantly, we tested whether the semantic advantage in non-decision time differed across conditions by calculating pairwise feature type x condition interactions on the original time scale. These interactions were calculated as the difference in the semantic-perceptual effect between two conditions. The absolute semantic advantage was reliably larger in the *Prospective* than in the *Immediate* condition (mean interaction = −48 ms, 94% HDI [−83, −15], posterior probability of a negative interaction = .995). Similarly, a reliable interaction was observed between the *Prospective* and *Delayed* conditions (−41 ms, 94% HDI [−79, −4], posterior probability = .984). In contrast, the interaction between the *Delayed* and *Immediate* conditions was small and uncertain (−8 ms, 94% HDI [−39, 28]). These findings indicate that the absolute semantic advantage in non-decision time was particularly pronounced when working memory representations had to be maintained in the face of interference, rather than following a longer retention interval alone. However, because non-decision time was modelled using a logarithmic link function, the corresponding interactions on the proportional scale were not reliably different from zero (semantic x Prospective: −0.038, 94% HDI [−0.120, 0.039]; semantic x Delayed: 0.026, [−0.058, 0.105]). Thus, although the interactions were supported on the original time scale, the larger absolute semantic advantage in Prospective trials may partly reflect longer overall non-decision times in this condition rather than a selective increase in the proportional efficiency of semantic processing.

We also observed consistent semantic advantages in drift rate across all three conditions (Fig. 6b). Semantic judgements were associated with higher drift rates than perceptual judgements in the *Immediate* condition (mean difference = 0.230, 94% HDI [0.130, 0.325]), the *Prospective* condition (0.241, [0.155, 0.327]) and the *Delayed* condition (0.265, [0.164, 0.359]). All three highest-density intervals excluded zero, indicating that semantic information contributed more efficiently to evidence accumulation irrespective of experimental condition. Direct comparisons revealed no reliable differences in the magnitude of this advantage between conditions (*Prospective* minus *Immediate*: 0.012, 94% HDI [−0.109, 0.143]; *Delayed* minus *Immediate*: 0.035, [−0.097, 0.172]; *Prospective* minus *Delayed*: −0.024, [−0.157, 0.104]). Thus, although semantic judgements consistently benefited from more efficient evidence accumulation, neither prospective interference nor increased retention time reliably modulated this advantage.

Finally, we examined whether semantic and perceptual judgements differed in decision thresholds (Fig. 6c). No reliable differences were observed in the *Immediate* condition (mean difference = −0.020, 94% HDI [−0.067, 0.029]), the *Prospective* condition (0.036, [−0.009, 0.081]) or the *Delayed* condition (0.039, [−0.010, 0.087]). Although the posterior estimates suggested numerically higher thresholds for semantic judgements in the Prospective and Delayed conditions, all intervals overlapped zero. Similarly, comparisons between conditions provided no reliable evidence that the semantic-perceptual difference in decision threshold varied with interference or retention time.

Together, these results indicate that semantic prioritisation in Experiment 2 is expressed through both shorter non-decision times and more efficient evidence accumulation. Semantic judgements consistently benefited from these two components across all experimental conditions, while decision-threshold differences were not reliably supported. The absolute semantic advantage in non-decision time was greatest in the Prospective condition, consistent with a potentially greater contribution of processes preceding evidence accumulation when working memory representations must be maintained in the face of interference. Semantic prioritisation appears to reflect a combination of advantages in processes outside evidence accumulation, potentially including retrieval or access to stored information, and the efficiency with which semantic information contributes to decision formation.

## Discussion

Across a reanalysis of our previous dataset and two new experiments, we found that categorical semantic judgements were consistently associated with shorter non-decision times than perceptual-format judgements, alongside advantages in evidence accumulation that varied across experimental contexts. Importantly, the relative contributions of these decision components were sensitive to attentional prioritisation. In Experiment 1, advance cueing of the relevant memory item substantially attenuated the semantic advantage in non-decision time, whereas the advantage in drift rate remained largely unchanged. Experiment 2 further demonstrated that semantic advantages in both non-decision time and drift rate were present across immediate retrieval, extended maintenance and intervening interference. Together, these findings suggest that semantic prioritisation in working memory cannot be attributed to a single decision process. Rather, the relative contributions of processes outside evidence accumulation, evidence accumulation itself and decision thresholds depend on the demands placed on working-memory representations.

Reanalysis of our earlier dataset using drift-diffusion modelling revealed a robust reduction in non-decision time for semantic judgements across memory loads and lags, consistent with semantic information influencing processes outside evidence accumulation following probe onset. This advantage was evident even when only a single item was maintained, whereas higher drift rates for semantic judgements were supported primarily at higher loads and longer lags. Allowing decision thresholds to vary further revealed differences in response caution under some conditions, suggesting that semantic prioritisation can reflect multiple, potentially opposing contributions to behavioural performance. These findings suggest a functional distinction between processes governing the timing with which information becomes usable for a decision and those governing the efficiency of evidence accumulation. Notably, this pattern resembles distinctions drawn in the episodic memory literature, where categorical semantic or conceptual information often precedes categorical perceptual information during retrieval, consistent with top-down or reconstructive access dynamics rather than feedforward perceptual processing (Lifanov et al., 2021; Linde-Domingo et al., 2019). However, reduced non-decision time cannot distinguish whether semantic information becomes available earlier through memory retrieval or through other probe- and decision-related processes. Moreover, because memory load and lag were analysed separately, their apparent modulation of drift- rate differences remains descriptive rather than evidence for formal interactions.

Experiment 1 provided a more direct test of whether semantic prioritisation depends on the ability to select and prepare a relevant memory representation before the probe. Behaviourally, semantic prioritisation was greater when participants could not pre-select a single item and instead had to maintain multiple representations concurrently. Drift-diffusion modelling revealed that under neutral cueing, semantic judgements showed substantially shorter non-decision times than perceptual judgements, whereas this difference was almost entirely eliminated when a valid retro-cue identified the relevant item in advance. In contrast, semantic judgements were associated with higher drift rates in both cueing conditions, with no reliable evidence that this advantage differed according to cue validity. Thus, attentional prioritisation selectively reduced the semantic-perceptual difference in non-decision time without a corresponding change in the relative efficiency of evidence accumulation. Whereas previous work has demonstrated that retro-cueing can preferentially enhance perceptual over semantic information (Cipriani et al., 2024), the present findings suggest that attentional prioritisation also influences how different dimensions of remembered information contribute to processes outside evidence accumulation. This dissociation indicates that attentional selection and subsequent decision formation are not interchangeable processes, and that the semantic advantage depends partly on when the relevant memory item can be prioritised.

One possible interpretation is that valid retro-cues allow the relevant memory representation to be selected or reactivated before the probe, reducing the processing demands that would otherwise have to be resolved following probe onset. Under neutral cueing, the relevant item cannot be identified in advance, potentially increasing the demands associated with selecting and reinstating task-relevant information. The attenuation of the semantic non-decision-time advantage under valid cueing is therefore consistent with semantic information being particularly useful when such processes must be completed at test. Importantly, the semantic and perceptual probe categories remained the same across cueing conditions, making a fixed difference in question processing alone insufficient to explain the interaction. Nevertheless, valid cueing may also influence preparatory processes, response strategies or expectations concerning the upcoming decision. The present results therefore implicate attentional prioritisation in processes outside evidence accumulation without uniquely identifying memory reactivation as the underlying mechanism.

These findings complement a growing literature demonstrating that semantic knowledge contributes to visual working-memory performance. Previous studies have shown that semantic associations and category coherence can facilitate the encoding and maintenance of multiple objects (Hu & Jacobs, 2021; O’Donnell et al., 2018; Yu et al., 2025), while meaningfulness can support the binding of object identity to spatial location (Sahar et al., 2024). These studies suggest that semantic knowledge can influence the organisation and stability of working-memory representations. The present findings address a related but distinct question of how judgements about categorical semantic information, relative to perceptual-format information, are expressed through different components of decision formation. In particular, the dissociation observed in Experiment 1 suggests that attentional prioritisation can substantially alter the semantic advantage in non-decision time while leaving the relative drift-rate advantage largely unaffected. However, these results do not establish whether semantic information was represented more faithfully, reactivated more efficiently or processed more readily once the relevant probe appeared.

In contrast to our previous findings (Kerrén et al., 2022), we did not observe a consistent relationship between semantic prioritisation and lag when attentional demands were controlled. In our earlier study, lag was necessarily confounded with memory load and serial position, such that increasing temporal distance also changed the circumstances under which individual items were maintained. Although Experiment 1 revealed lag-related increases in semantic prioritisation in the valid condition, most clearly in accuracy and the balanced integration score, this relationship was not observed under neutral cueing. When memory load was held constant and retention interval was manipulated independently of interference in Experiment 2, temporal delay alone did not reliably increase the relative semantic advantage. Together, these findings argue against a simple account in which the prioritisation of semantic information necessarily increases with the passage of time over the intervals examined.

One possible explanation comes from work by Oberauer (Oberauer, 2002), which distinguishes information within the focus of attention from other information maintained in working memory. Under this framework, representations may remain available without occupying the focus of attention continuously. Extending a retention interval therefore does not need to produce a corresponding increase in the demands associated with accessing a representation at test. In our Delayed condition, participants could maintain the studied items without performing an additional task, potentially allowing information to remain available without substantial competition from concurrent processing. This interpretation is consistent with accounts proposing that information outside the focus of attention can remain available for subsequent use, and that temporal delay does not need to be equivalent to attentional disruption (Oberauer, 2002; Oberauer & Hein, 2012).

Experiment 2 further dissociated the effects of temporal delay from those of an intervening task during maintenance. Semantic judgements were associated with both shorter non- decision times and higher drift rates across Immediate, Delayed and Prospective conditions. The absolute semantic-perceptual difference in non-decision time was larger in the Prospective condition than in either the Immediate or Delayed condition, consistent with a greater absolute advantage when working-memory representations had to be maintained while participants performed a competing task. However, these interactions were supported on the original time scale but not on the logarithmic scale used to estimate the model. The larger difference in milliseconds may therefore partly reflect longer overall non-decision times in Prospective trials, rather than a selective increase in the proportional semantic advantage. Furthermore, the semantic drift-rate advantage was comparable across all three conditions, suggesting that interference did not reliably alter the relative efficiency with which semantic and perceptual information contributed to evidence accumulation.

These findings are relevant to accounts emphasising interference, rather than the mere passage of time, as a determinant of working-memory performance (Oberauer & Lin, 2017; Portrat et al., 2008; Ricker et al., 2010). Although the behavioural and modelling results are consistent with interference placing additional demands on the processing of remembered information, they do not establish that such demands selectively enhance semantic retrieval. The Prospective condition also required participants to perform an intervening decision and motor response before the memory probe, introducing task-switching and response- preparation demands that cannot be fully separated from interference with maintained representations. The results therefore suggest that attentional disruption and retention time can have different consequences for decision formation, but do not uniquely identify representational protection or reinstatement as the source of the larger absolute semantic advantage.

Taken together, these findings are compatible with theories distinguishing a narrow focus of attention from a broader set of activated but unattended working-memory representations (Cowan, 2001; Oberauer, 2002; Oberauer & Lin, 2017) as well as accounts proposing that unattended or activity-silent working memory may rely on mechanisms overlapping with episodic memory retrieval (Beukers et al., 2021; Rose et al., 2016; Stokes, 2015). Within such frameworks, information outside the focus of attention may require additional processing before it can guide behaviour. The present findings suggest that categorical semantic and perceptual-format information differ in how they contribute to these processes, particularly when advance attentional prioritisation is unavailable. One possibility is that semantic information remains relatively accessible when memory representations are not actively prioritised, potentially because categorical information can be reconstructed or used for decision-making more readily than perceptual-format information. However, the present data do not establish that unattended representations are transformed into more abstract or long- term-memory-like formats.

This distinction is important when relating the present findings to theories of working-memory representation. Neural decoding studies have demonstrated that perceptual information can be maintained in sensory cortical regions (Harrison & Tong, 2009; Serences et al., 2009), while other theoretical accounts distinguish different representational states or mechanisms of maintenance (Beukers et al., 2021; Rose et al., 2016; Stokes, 2015). Such approaches provide evidence concerning the content and maintenance of working-memory representations that is complementary to the present decision-model analysis. Similarly, models emphasising constraints on the precision or resolution of stored representations (Zhang & Luck, 2008) address a different level of explanation from the decision components considered here. Drift-diffusion parameters characterise how choices and reaction times arise following a probe; they do not directly measure representational content, fidelity or format. Consequently, differences in non-decision time or drift rate cannot themselves demonstrate that perceptual representations are replaced by semantic ones, or that unattended working memory relies on long-term-memory-like representations. Establishing such transformations would require direct representational measures.

A related possibility is that semantic prioritisation reflects more general differences in the processes through which internally maintained information becomes available for use. During perception, visuoperceptual attributes can become available before conceptual properties, whereas previous work on episodic retrieval has demonstrated the opposite ordering for comparable categorical judgements (Lifanov et al., 2021; Linde-Domingo et al., 2019). Our previous findings using the same stimulus space suggested that a similar ordering may characterise visual working memory (Kerrén et al., 2022). The present results extend this comparison by showing that semantic prioritisation can involve both processes outside evidence accumulation and the accumulation of decision evidence, with attentional prioritisation exerting a particularly pronounced influence on the former. This pattern is compatible with theories proposing that internally generated representations are accessed through reconstructive or reactivation-related processes, as discussed in the literature on activity-silent working memory, episodic retrieval and visual imagery (Dijkstra et al., 2018, 2020; Rose et al., 2016; Stokes, 2015). Nevertheless, the correspondence between working- memory and episodic-retrieval dynamics remains a functional analogy rather than evidence that the same neural or computational mechanisms are responsible.

An important limitation concerns the operationalisation of semantic and perceptual information. Here, semantic judgements involved a binary animacy-related category distinction, whereas perceptual judgements concerned the format in which an object was presented. Although this design permitted a controlled comparison between two task-relevant dimensions, both judgements were categorical, and they may have differed in the ease with which the relevant information could be evaluated or mapped onto a response. Moreover, semantic knowledge encompasses considerably richer structures, including graded conceptual similarity, functional relationships and contextual associations. Recent evidence suggests that categorical, semantic and visual similarity can make distinct contributions to short-term and long-term memory search (Shang et al., 2026). Consequently, the present findings should not be interpreted as establishing a general processing advantage for all forms of semantic information. Rather, they identify differences between categorical semantic and perceptual-format judgements under specific working-memory demands. Future work should examine whether similar decision-stage effects generalise to richer semantic relationships, graded perceptual properties and other combinations of stimulus dimensions.

A further limitation concerns the interpretation of non-decision time. This parameter encompasses processes outside evidence accumulation, including probe encoding, preparatory processing, retrieval-related operations and motor execution. The observed semantic advantage therefore cannot be attributed uniquely to faster retrieval of stored representations. Although the selective attenuation of this advantage under valid cueing constrains accounts based solely on fixed probe difficulty, it does not eliminate explanations involving differences in response preparation, expectations or decision strategy. Future experiments could independently manipulate information about the relevant memory item and the forthcoming response, allowing memory selection and response preparation to be separated more directly. Combining such manipulations with time-resolved neural measures could further establish when categorical semantic and perceptual information becomes available relative to the onset of evidence accumulation.

More broadly, our findings suggest that understanding working-memory limitations requires consideration not only of the amount or fidelity of retained information, but also of the processes through which different aspects of that information are used to guide decisions. Semantic information was associated with shorter non-decision times across a range of working-memory demands and, in many conditions, with more efficient evidence accumulation. Crucially, advance attentional prioritisation selectively reduced the former advantage without reliably changing the latter. This dissociation provides evidence that attentional selection influences how remembered information contributes to subsequent decisions, while remaining compatible with a possible role for retrieval-related or reconstructive processing. Whether these decision-stage differences reflect changes in representational format, retrieval dynamics or other preparatory mechanisms remains an important question for future research.

## Method

### Reanalysis of Kerrén et al. 2022

We reanalysed data from Kerrén et al. (2022), in which participants (n = 103 after exclusions) performed a visual working memory task requiring maintenance of one to four object images across a short delay. Each object varied orthogonally in semantic category (animate/inanimate) and visuo-perceptual format (photograph/drawing). At the test, participants answered either a semantic or a perceptual question about one studied item, where reaction time and accuracy were obtained. Memory load and the lag between encoding and test were systematically varied. The present analyses focus on Experiment 2, which included both load and lag manipulations. Load was defined by how many items were presented during the study phase. Lag was defined by the distance between study and test. For a trial of three items, the first item studied would have the largest lag, whereas the last item studied would have the shortest lag. Full procedural details are reported in the original study. Here, we applied hierarchical drift diffusion modelling to dissociate pre-decisional access from evidence accumulation processes.

### Experiment 1

#### Participants

We recruited 96 participants online (48 female, 47 male, and 1 did not prefer to say) with a mean age of 29.69 years (SD ± 4.18 years, range = 21-35 years). Participation was with informed consent via the Prolific platform (https://www.prolific.ac/). The eligibility criteria were that participants had to be between 18 and 35 years old, fluent in English, being from the United Kingdom or the United States, normal, or corrected-to- normal vision, have a minimum of 30 previous survey submissions, and have a minimum approval rate of 95 on Prolific. Participants were reimbursed with £4.5 for taking part in the experiment. The experiment lasted approximately 30 minutes. The experiment was terminated prematurely for participants who failed to pass attention checks (see below), which resulted in no payment (n = 0 participants). We also excluded participants whose accuracy did not significantly exceed chance level (p<.05, Binomial test against 50% correct responses) in either the semantic or perceptual judgments (n = 15 participants). Thus, 81 participants remained for analysis. The experiment was approved by the University of Granada Ethics Committee. Sample size and exclusion criteria was pre-registered before data collection (see https://researchbox.org/5966).

#### Stimuli

The experimental stimuli comprised 150 pictures (75 depicting animals, 75 depicting inanimate objects). An additional 10 pictures were used for instructions and practice trials. A detailed description of the stimulus set is provided in (Linde-Domingo et al., 2019). For each of the pictures (which are colour photographs), a line-drawing version (black-and-white) was created using GNU image manipulation software (http://www.gimp.org). For each participant, half of the animals and half of the inanimate objects were shown as line drawings (randomly assigned) and the others as colour photographs. The stimuli thus differed in two orthogonal dimensions, visuo-perceptual (photograph/drawing) and semantic (animate/inanimate; Fig. 1a).

#### Task

Each trial started with a central fixation cross (500ms), after which 3 pictures were sequentially presented (1500ms each) at different random locations on the screen (Fig. 1b). The locations were randomly selected from 16 equidistant positions on an invisible circle around screen centre. The positions were selected to be at minimum 45° apart from each other. Participants were asked to remember the pictures and locations. Pictures were chosen so that within a trial they differed on experimental features. That is, if we imagine this being in a trial of two items, one would always be an animal and one would always be an object. Similarly, one of them would be line-drawing and the other a photograph. Naturally, this is not possible in a trial of three items, where we instead counterbalanced across trials, such that two coloured items were paired with a line-drawn item equal number of times as the opposite. And the same for animals and objects. After a 1000ms blank screen, a retro-cue was shown for 500ms. In the *valid cue* condition, a circle was shown in the location where the questions would later appear. In the *neutral cue* condition, three circles were shown in the three locations that items were previously studied. After a delay (1000ms), a question appeared at the location of one of the objects. The question was aimed at either probing the semantic (“Animal or Object?”) or perceptual (“Photo or Drawing?”) features of the studied item that was presented at that location. These questions were counterbalanced within participants such that the semantic and perceptual questions were equally likely to be asked first. In addition, across participants we also counterbalanced the order of words in the question, (e.g., “Object or Animal?” and “Drawing or Photo?”) and how these different options were tested together (e.g., if “Object or Animal?” and “Drawing or Photo?” were the two response options for one participant, then “Animal or Object?” and “Drawing or Photo?” could be the response options for the next participant and so on). Participants were asked to select the response option to report its characteristics (perceptual and semantic) using the right and left arrow keys. Participants were given 3 seconds to make their selections (where they could not click a button during the first 200ms). Upon selection the font colour of the question turned red (for incorrect) or green (for correct). Independent if participants responded or not within 3 seconds of the first question, the next question appeared (if the first one probed semantic features, the second probed perceptual features and vice versa). Again, independent of engagement from participants, a new trial began after 3 seconds. All trials with no response were excluded from analysis.

#### Procedure

After extensive written instructions, participants performed three practice trials to familiarise themselves with the task. Participants were free to repeat the instructions and practice trials until they felt confident at performing the task. Thereafter, each participant performed 100 experimental trials (50 in each condition). After the 4th, 16th, and 28th trial, a brief attention check was performed. For this, a large filled triangle was presented centrally in either green, blue, or red colour and participants were asked to name the colour via button press. If a participant failed on any of the attention checks, the experiment was aborted (see above, *Participants*). After 50 trials, participants could take a short break (self-paced).

### Experiment 2

#### Participants

We recruited 110 participants online (64 female, 45 male, and 1 who did not prefer to say) with a mean age of 29.87 years (SD ± 4.28 years, range = 20-35 years). The task took approximately 35 minutes and participants were compensated £6 for taking part in the experiment. Again, we used attention checks, and the experiment was terminated prematurely for participants who failed to pass attention checks, which resulted in no payment (n = 1 participants). We also excluded participants whose accuracy did not significantly exceed chance level (p<.05, Binomial test against 50% correct responses) in either the semantic or perceptual judgments (n = 33 participants). Thus, 77 participants remained for analysis. Sample size and exclusion criteria was pre-registered before data collection (see https://researchbox.org/5966). The experiment was approved by the University of Granada Ethics Committee.

#### Stimuli, Task, and Procedure

The design of Exp. 2 was similar to Exp. 1, with the following differences. After having studied 3 items, at test, a cue signalling which condition they were in, was shown for 500ms with a delay of 1000ms both preceding and following the condition- cue. The cue was “Now”, “Later” or “Wait”, depending on the condition they were in. If the cue was “Now” (*Immediate* condition) it meant that the upcoming questions would be administered immediately following the delay (equal to the *neutral* condition in Exp. 1). If the cue was “Later” (*Prospective* condition), participants were first exposed to a symbol (triangle, square or circle, randomised across trials) and a question (triangle?, square?, or circle?, randomised across trials), and the response options (yes / no, or no / yes, counterbalanced). They had 2000ms to respond, whereafter the questions regarding the studied items appeared. If the cue was “Wait” (*Delayed* condition), a symbol made up of stacking the triangle, circle and square on top of each other was shown for 2000ms. The task for the participants was to just observe the symbol, whereafter the questions regarding the studied items appeared. To equate cognitive load in the three conditions, in the immediate condition, the questions regarding the symbols appeared after participants had responded to the questions regarding the studied items. Participants performed a total of 100 trials, divided into 34, 33, and 33 trials in each condition.

### Data analysis

The BIS was calculated by standardising (z-scoring) the RTs on correct trials and the accuracies relative to their mean and standard deviation over all conditions, and then subtracting within each condition the RT score from the accuracy score (Liesefeld et al., 2015). Main analyses were performed in MATLAB, version 2022b (The MathWorks, Munich, Germany) and Bayesian ANOVAs were performed on JASP 0.95.4 using default priors and computing effects across matched models.

#### Drift diffusion modelling

We fitted hierarchical drift diffusion models (HDDMs) to single-trial choice and reaction time data using the *HSSM package* in Python (Fengler et al., 2025). Analyses were restricted to reaction times greater than 200ms, corresponding to the minimum response interval enforced by the experimental task. Responses were coded as correct (+1) or incorrect (−1), and the principal predictor was Feature, distinguishing semantic (animate versus object) from perceptual (photograph versus drawing) judgements. To accommodate occasional lapses and responses not adequately described by the diffusion process, we used HSSM’s default mixture-model approach, specifying a fixed outlier probability of 0.05. Under this specification, each trial contributes to a mixture of the DDM likelihood and a uniform lapse distribution, rather than identifying and excluding a fixed proportion of trials.

The models distinguished three components of decision-making: non-decision time (*t*), reflecting processes occurring outside evidence accumulation, including stimulus encoding, preparatory processing and motor execution; drift rate (*v*), reflecting the rate and efficiency of evidence accumulation towards a response; and decision threshold (*a*), reflecting the amount of evidence required before a decision is made. These parameters were estimated within a hierarchical Bayesian framework, with participant-specific random intercepts allowing individual differences while partially pooling information across participants. Starting-point bias (*z*) was estimated as a shared parameter but was not allowed to vary with experimental predictors.

To improve sampling efficiency and ensure that estimated parameters remained within their permissible ranges, we used HSSM’s *log_logit* link-function specification, applying logarithmic links to non-decision time and decision threshold while retaining an identity link for drift rate. We used the package’s *safe prior* specification, with additional weakly informative normal priors on the intercepts of drift rate (Normal[0, 0.5]) and log-transformed non-decision time (Normal[log(0.2), 0.5]). Remaining regression coefficients and hierarchical variance parameters were assigned HSSM’s weakly informative priors. Consequently, coefficients for non-decision time and decision threshold were estimated on a logarithmic scale, whereas drift- rate coefficients were estimated on their original scale.

Models were estimated using the No-U-Turn Sampler (NUTS) implemented through the NumPyro backend. For the final models, we ran four independent chains, each with 3000 tuning iterations followed by 3000 retained posterior draws, yielding 12000 posterior samples per model. The target acceptance probability was set to 0.95. One model in the reanalysis of Kerrén et al. (2022), corresponding to memory load 1, was additionally refitted using 5000 tuning iterations and 4000 retained draws per chain to improve convergence. Convergence was assessed using the rank-normalised potential scale reduction statistic (R-hat), bulk and tail effective sample sizes, and the number of divergent transitions. Posterior distributions were summarised using posterior means and 94% highest-density intervals (HDIs).

Sampling convergence was assessed using R-hat, bulk and tail effective sample sizes (ESS), and divergent transitions (Supplementary Table 1). No divergent transitions were observed. Convergence was satisfactory for the population-level experimental coefficients, although some participant-specific parameters in the load 1 reanalysis showed elevated R-hat values and reduced effective sample sizes.

#### Model specification and comparison

We initially evaluated four model specifications to determine which components of decision- making contributed to semantic prioritisation.

*Model 1 (non-decision time)* allowed Feature to modulate non-decision time (*t*), with participant-specific random intercepts. Drift rate and decision threshold were not allowed to vary with Feature.

*Model 2 (drift rate)* allowed Feature to modulate drift rate (*v*), with participant-specific random intercepts, while non-decision time and decision threshold were not allowed to vary with Feature.

*Model 3 (non-decision time and drift rate)* allowed Feature to modulate both *t* and *v* jointly, each with participant-specific random intercepts, while decision threshold was not allowed to vary with Feature.

*Model 4 (non-decision time, drift rate and decision threshold)* extended Model 3 by additionally allowing Feature to modulate decision threshold (*a*), with participant-specific random intercepts for all three parameters. This model therefore allowed semantic–perceptual differences to arise independently or jointly through processes outside evidence accumulation, the efficiency of evidence accumulation, and the amount of evidence required before responding.

Initial model comparisons were performed using Pareto-smoothed importance sampling leave-one-out cross-validation (PSIS-LOO), implemented in *ArviZ*. However, substantial Pareto-*k* diagnostic violations indicated that the resulting predictive estimates were unreliable. We therefore evaluated the predictive performance of Models 3 and 4 using five-fold held-out cross-validation. Trials were divided into five folds, stratified within participants and, where possible, by experimental condition and feature type. Each model was fitted to four folds and evaluated on the remaining fold, ensuring that training and test sets contained trials from the same participants. Held-out log predictive densities were calculated by averaging predictive likelihoods across posterior draws and summed across folds to obtain the expected log predictive density (ELPD). Differences in predictive performance were evaluated using paired held-out observations, with uncertainty estimated from 10000 participant-level bootstrap resamples to account for the dependence between trials from the same participant.

Five-fold cross-validation consistently favoured Model 4 over Model 3. In Experiment 1, Model 4 provided higher predictive accuracy (ΔELPD = 107.92, participant-bootstrap 95% CI [64.06, 155.44]). The same preference was observed in the reanalysis of Kerrén et al. (2022), both when grouping trials by study-test lag (lag 3: ΔELPD = 20.86, 95% CI [8.05, 34.87]) and by memory load (load 3: ΔELPD = 29.61, 95% CI [1.99, 59.84]). Based on these comparisons, we adopted Model 4 as the common specification for the final analyses.

#### Experiment-specific analyses

For the reanalysis of Kerrén et al. (2022), models were fitted separately for memory loads 1- 4 and lags 1-4. For each load and lag, Feature was included as a predictor of the relevant DDM parameters. The lag analyses used the temporal distance between the presentation of the probed item and the subsequent test, with lag 1 indicating the shortest and lag 4 the longest study-test interval. The same three-parameter model was applied across all load and lag conditions to ensure comparability of parameter estimates.

For Experiment 1, Feature, cue validity (valid versus neutral) and their interaction were included as predictors of non-decision time, drift rate and decision threshold. This allowed us to examine whether semantic-perceptual differences in each decision component depended on the availability of an informative attentional cue.

For Experiment 2, we directly adopted Model 4 based on its superior predictive performance in the preceding analyses, without repeating model selection. Feature, experimental condition (Immediate, Prospective, Delayed) and their interaction were included as predictors of all three DDM parameters. The Immediate condition served as the reference level, enabling direct estimation of the semantic–perceptual differences in each condition and comparisons of these effects across conditions.

#### Posterior contrasts and statistical inference

To quantify semantic prioritisation, we calculated posterior contrasts between semantic and perceptual judgements for each parameter and experimental condition. For non-decision time and decision threshold, posterior estimates were transformed from the logarithmic scale to their original parameter scales before calculating contrasts. Non-decision-time differences were expressed in milliseconds, whereas drift-rate and threshold differences were reported in their respective model units.

For Experiment 1 and Experiment 2, interactions were additionally examined by comparing the semantic-perceptual contrasts between experimental conditions. These were calculated as differences of posterior differences, preserving the correspondence between posterior draws. For non-decision time, this provided interaction estimates on the original time scale, reflecting differences in the absolute semantic advantage between conditions. Because logarithmic link functions were used, we additionally examined the interaction coefficients on the model’s logarithmic scale, which quantify changes in proportional rather than absolute effects.

Statistical inference was based on the posterior distributions of these contrasts, with 94% HDIs and posterior probabilities used to characterise the direction and uncertainty of each effect. Intervals excluding zero were interpreted as providing evidence for a directional difference.

For the separately fitted load- and lag-specific models, differences in effect magnitude across conditions were treated descriptively rather than as formal tests of interactions.

## Data availability

The data that support the findings of this study are available at https://github.com/kerrencasper/Flipper_2025_git

## Code availability

The code to reproduce the results of this study is available at https://github.com/kerrencasper/Flipper_2025_git

## Acknowledgments

C.K. is supported by the Max Planck society. C.G.G. is supported by Project PID2023- 149428NB-I00 funded by MCIN/AEI/10.13039/501100011033 and by FEDER, EU, and Grant RYC2021-033536-I funded by MCIN/AEI/10.13039/501100011033 and by the European Union NextGeneration EU/PRTR. J.L.D. is supported by Project PID2023-151104NA-I00 funded by MCIN/AEI/10.13039/501100011033 and by FEDER, EU, and Grant RYC2021- 033940-I funded by MCIN/AEI/10.13039/501100011033 and by the European Union NextGeneration EU/PRTR.The Mind, Brain and Behavior Research Center receives funding from grants CEX2023-001312-M by MICIU/AEI/10.13039/501100011033 and UCE-PP2023-11 by the University of Granada. This work was supported by the Unidad de Excelencia María de Maeztu – CIMCYC (CEX2023-001312-M), funded by MICIU/AEI/10.13039/501100011033.

The funders had no role in the study design, data collection and analysis, decision to publish or preparation of the manuscript.

## Author contributions

C.K., J.L.D. and C.G.G. were responsible for conceptualisation, project administration, resources, data collection, data curation, and formal analysis. All authors were involved in methodology, visualisation, and writing the original draft, reviewing, and editing.

## Competing interests

The authors declare no competing interests.

## Supplemental material

**Supplementary Figure 1.**
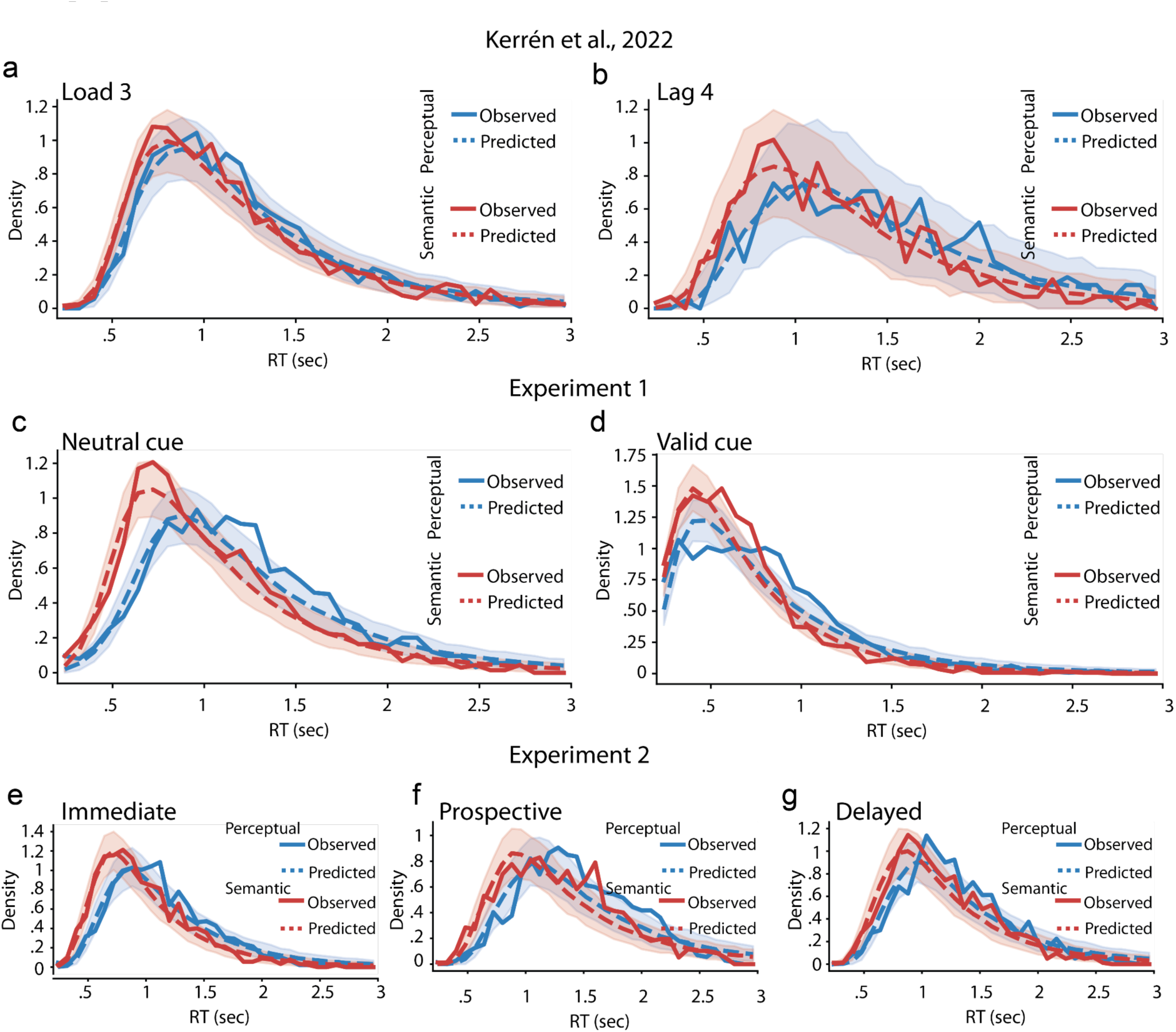
Posterior predictive checks of the hierarchical drift-diffusion models. Observed and model-predicted reaction-time distributions for correct semantic and perceptual judgements, obtained from the final Model 4 drift-diffusion model. **a-b** Reanalysis of Kerrén et al. (2022), showing memory load 3 and lag 4, respectively. **c-d** Experiment 1, showing neutral and valid retro-cue conditions, respectively. **e-g** Experiment 2, showing Immediate, Prospective and Delayed conditions, respectively. Solid lines represent observed reaction-time distributions, whereas dashed lines represent the mean posterior predictive distributions. Shaded areas indicate the central 94% posterior predictive intervals. Blue and red indicate perceptual and semantic judgements, respectively. Across datasets and experimental conditions, the model broadly reproduced the observed reaction-time distributions and the differences between semantic and perceptual judgements.

**Supplementary Table 1.** Convergence diagnostics for the final hierarchical drift- diffusion models.

| Model | Chains | Draws/chain | Max R-hat | Min bulk ESS | Min tail ESS | Divergences | R-hat > 1.01 |
|---|---|---|---|---|---|---|---|
| Experiment 1 | 4 | 3000 | 1.01 | 545 | 607 | 0 | 0 |
| Experiment 2 | 4 | 3000 | 1.00 | 670 | 789 | 0 | 0 |
| Kerrén Load 1 | 4 | 3000 | 1.10 | 27 | 183 | 0 | 8 |
| Kerrén Load 2 | 4 | 3000 | 1.00 | 993 | 759 | 0 | 0 |
| Kerrén Load 3 | 4 | 3000 | 1.01 | 817 | 289 | 0 | 0 |
| Kerrén Load 4 | 4 | 3000 | 1.00 | 855 | 512 | 0 | 0 |
| Kerrén Lag 1 | 4 | 3000 | 1.00 | 2324 | 2528 | 0 | 0 |
| Kerrén Lag 2 | 4 | 3000 | 1.01 | 730 | 1012 | 0 | 0 |
| Kerrén Lag 3 | 4 | 3000 | 1.01 | 644 | 994 | 0 | 0 |
| Kerrén Lag 4 | 4 | 3000 | 1.01 | 203 | 507 | 0 | 0 |

## Notes

### Competing Interest Statement

The authors have declared no competing interest.

### Summary of Updates

In light of reviewers' comments, we have substantially updated the manuscript.

